# Donor Strand Complementation and Calcium Ion Coordination Drive the Chaperone-free Polymerization of Archaeal Cannulae

**DOI:** 10.1101/2024.12.30.630787

**Authors:** Mike Sleutel, Ravi R. Sonani, Jessalyn G. Miller, Fengbin Wang, Andres Gonzalez Socorro, Yang Chen, Reece Martin, Borries Demeler, Michael J. Rudolph, Vikram Alva, Han Remaut, Edward H. Egelman, Vincent P. Conticello

**Affiliations:** Structural Biology Brussels, Vrije Universiteit Brussel, Brussels, Belgium; Structural and Molecular Microbiology, VIB-VUB Center for Structural Biology, Brussels, Belgium; Department of Biochemistry and Molecular Genetics, University of Virginia, Charlottesville, VA, 22908, USA; Department of Chemistry, Emory University, Atlanta, GA, 30322, USA; New York Structural Biology Center, 89 Convent Avenue New York, NY, 10027; Biochemistry and Molecular Genetics Department, University of Alabama at Birmingham, Birmingham, AL, 35233, USA; Department of Chemistry and Biochemistry, University of Lethbridge, Lethbridge, Alberta T1K 3M4, Canada; Department of Protein Evolution, Max Planck Institute for Developmental Biology Tübingen, Tübingen 72076, Germany; The Robert P. Apkarian Integrated Electron Microscopy Core (IEMC), Emory University, Atlanta, GA, 30322, USA

## Abstract

Cannulae are tubular protein filaments that accumulate on the extracellular surface of the hyperthermophilic archaeon *Pyrodictium abyssi* during cell division. Cannulae have been postulated to act as a primitive extracellular matrix through which cells could communicate or exchange material, although their native biological function remains obscure. Here, we report cryoEM structural analyses of *ex vivo* cannulae and of *in vitro* protein assemblies derived from recombinant cannula-like proteins. Three-dimensional reconstructions of *P. abyssi* cannulae revealed that the structural interactions between protomers in the native and recombinant filaments were based on donor strand complementation, a form of non-covalent polymerization in which a donor β-strand from one subunit is inserted into an acceptor groove in a β-sheet of a neighboring subunit. Donor strand complementation in cannulae is reinforced through calcium ion coordination at the interfaces between structural subunits in the respective assemblies. While donor strand complementation occurs during the assembly of chaperone-usher pili, this process requires the participation of accessory proteins that are localized in the outer membrane. In contrast, we demonstrate that calcium ions can induce assembly of cannulae in the absence of other co-factors. Crystallographic analysis of a recombinant cannula-like protein monomer provided evidence that calcium ion binding primes the precursor for donor strand invasion through unblocking of the acceptor groove. Bioinformatic analysis suggested that structurally homologous cannula-like proteins occurred within the genomes of other hyperthermophilic archaea and were encompassed within the TasA superfamily of biomatrix proteins. CryoEM structural analyses of tubular filaments derived from *in vitro* assembly of a recombinant cannula-like protein from an uncultured *Hyperthermus* species revealed a common mode of assembly to the *Pyrodictium* cannulae, in which donor strand complementation and calcium ion binding stabilized longitudinal and lateral assembly in tubular 2D sheets.

## Introduction

Filamentous protein assemblies are common in biological systems in which they are involved in a wide range of functions that are critical to the survival and propagation of the organism.^1^ Many of these functions, e.g., locomotion,^2^ adhesion,^3,4^ tunable mechanical response,^3–8^ multicellular organization,^9–12^ electrical conductivity,^13–16^ directional transport of substrates,^17–21^ regulation of enzymatic catalysis,^22^ etc., would be desirable to emulate in synthetic protein-based nanomaterials that could be tailored for specific applications.^23^ However, this approach requires prior knowledge of the sequence-structure-function relationships that are responsible for the observed behavior of the corresponding biologically derived assemblies, as well as convenient methods for the fabrication of the protein filaments. The physical principles that underlie the function of these biologically derived protein-based nanomaterials are being revealed through structural analysis at near-atomic resolution using electron cryomicroscopy (cryoEM).^24,25^ These studies provide insight into the relationship between structure and biological function that could provide a starting point for *de novo* design of synthetic analogues that mimic the function of native protein assemblies.^26^ High-resolution structural information can also enable an understanding of the molecular-level interactions that underlie the mechanism of self-assembly, thereby facilitating the development of reliable methods for recombinant production of the precursor proteins and their *in vitro* or *in vivo* conversion into functional filaments with desired properties.^27,28^

A scientifically and technologically interesting class of protein-based assemblies for the development of synthetic biomaterials are cannulae, which are tubular structures that form during the life cycle of the hyperthermophilic archaeon *Pyrodictium abyssi* and related organisms in the family *Pyrodictiaceae*.^29–32^ These protein-based filaments have been observed on the surface of *P. abyssi* and *occultum* strains under culture conditions that mimic the native growth environment.^31–33^ Members of the family *Pyrodictiaceae* are obligate anaerobes that propagate under hyperthermophilic deep-sea conditions with optimal growth rates in the temperature range from 80°C to 110 °C.^34^ The persistence of cannulae under these conditions suggested that the filaments display significant thermostability and mechanical stiffness, which has attracted biotechnological interest for potential use as robust delivery platforms for controlled release.^35^ However, *Pyrodictium* species are difficult to culture at preparative scale and not genetically tractable at present.^36^

The cannulae of *Pyrodictium abyssi* have been the most thoroughly studied experimental system. Cell cultures of *P. abyssi* displayed robust growth and achieved high cell densities albeit under restrictive culture conditions, e.g., high temperature and corrosive media.^32^ Dark field microscopy of growing cultures of *P. abyssi* provided evidence that the flat cells divided through binary fission.^36,37^ During cell division, cannulae are synthesized on the cell surface such that the daughter cells remain connected after separation. These extracellular connections were maintained in subsequent rounds of cell division, which resulted in formation of a dense network in which cells are linked through cannulae. Under cell culture, *P. abyssi* cells were never observed without cannulae, which have been postulated to serve as a primitive extracellular matrix through which cells could communicate or exchange material. Cryo-electron tomography (cryoET) studies of *P. abyssi* cultures^38^ indicated that the cannulae embedded in the pseudo-periplasmic space of inter-connected cells, which were typically covered with an S-layer. While the biological function of cannulae remains obscure, cryo-ET analysis provided evidence for density within the lumen of a cannula connecting *P. abyssi* cells, which led to a proposed role for cannulae in substrate transport. Here, we report cryoEM structural analyses of *ex vivo* cannulae and of *in vitro* protein assemblies derived from recombinant cannula-like proteins.^29–32^ Three-dimensional reconstructions of the *ex vivo* and *in vitro* cannulae from *P. abyssi* revealed that the structural interactions between protomers in both filaments were based on donor strand complementation, a form of non-covalent polymerization where a β-strand from one subunit is inserted into a β-sheet in a neighboring subunit, that has been observed in structures of presumably unrelated bacterial and archaeal pili.^3–7,9–11,39,40^ In cannulae, this interaction is reinforced through a network of coordination interactions between structural subunits that is mediated by calcium ion binding. Crystallographic analysis of a recombinant cannula-like protein monomer provided insight into the role of calcium ion on the self-assembly of the corresponding filament. Bioinformatic analysis suggested that structurally homologous cannula-like proteins occurred within the genomes of other hyperthermophilic archaea and were encompassed within the TasA superfamily of biomatrix proteins. CryoEM structural analysis of tubular filaments derived from *in vitro* assembly of a recombinant cannula-like protein from an uncultured *Hyperthermus* species revealed a common mode of assembly to the *Pyrodictium* cannulae, in which donor strand complementation and calcium ion binding stabilized longitudinal and lateral assembly in tubular 2D sheets. The facile *in vitro* assembly of cannula-mimetic tubules represents an attractive approach for the fabrication of thermodynamically stable and chemically addressable filamentous protein nanomaterials. In contrast to chaperone-usher pili, in which the protein subunits undergo polymerization with strand donation requiring both a chaperone and an usher, the recombinant cannula-mimetic proteins self-assemble spontaneously in the presence of calcium ions in aqueous solution under controlled conditions *in vitro* and maintain a similar structure to the *ex vivo P. abyssi* filament.

## Results

To gain structural insight into the cannulae fibers found on the surface of *Pyrodictium abyssi*, we initiated a cryoEM study of filaments observed from culture of *P. abyssi* AV2 for which the genomic sequence was available (NZ_AP028907.1). From the raw micrographs, two main fiber populations could be discerned under the culture conditions (see *Materials and Methods*), namely cannulae and an unidentified class of pili (see below). Cannulae were present as long (> 5 µm) filaments with a diameter of ∼26 nm that tended to cluster into large rafts (Supplementary Figure 1a). High magnification imaging (Figure 1d, Supplementary Figure 1b) revealed thinner, flexible (∼2 nm diameter) fibrils that were either associated to the cannulae or freely floated in suspension. Although these 2 nm fibrils were mostly seen to be randomly bound to cannulae filaments, examples were found in which the fibril as helically wound inside the cannulae scaffold seemingly following the helical pitch (Figure 1d), which was suggestive of a specific interaction between both fiber types.

**Figure 1:**
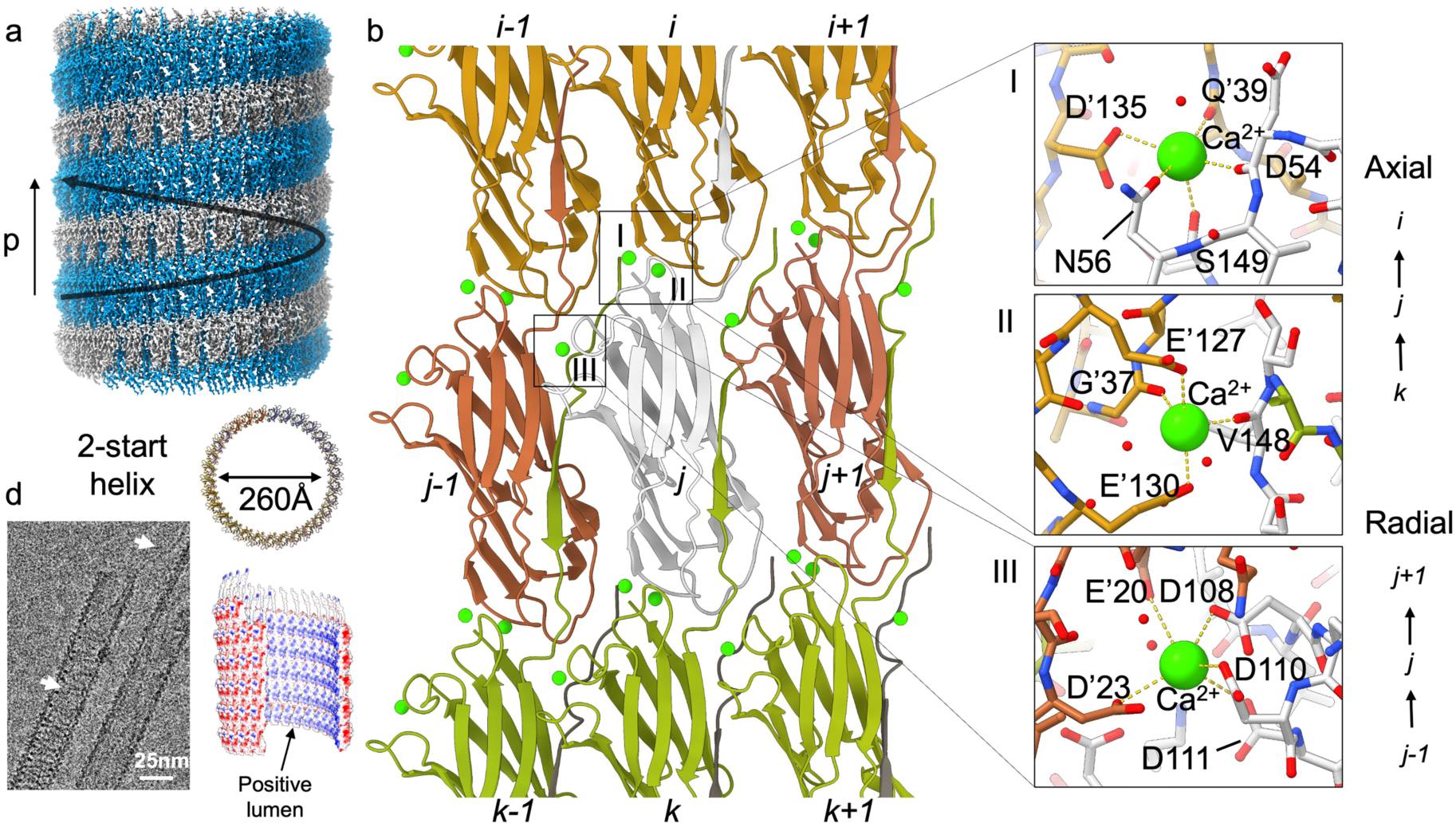
**CryoEM structure of *ex vivo* cannulae from the hyperthermophilic archaeon *Pyrodictium abyssi* AV2**. (a) cryoEM volume revealing a two-start (C2) helical fiber with a pitch *p* and diameter of 94.6 Å and 260 Å, and a rise and twist of 3.18 Å and 12.1°, respectively; (b) local interaction network between neighboring cannulae: subunits interlock through donor strand complementation along the axial (e.g. *i*→*j*→*k*) direction and partake in further intermolecular contacts mediated by calcium coordination along the axial and radial (e.g. *j*-1→*j*→*j*+1) direction; (c) stick representation of three types of calcium pockets: (i) axial stacking of subunits via type I and II calcium binding centers, (ii) radial contacts are mediated via type III Ca^2+^ binding sites; (d) raw cryoEM image of a cannulae filaments loaded with a helically winding thinner filament (2 nm; cargo). White arrows denote winding cargo and dangling cargo associated with the cannulae filaments; (e) surface electrostatics showing a predominantly negatively charged surface and a positively charged lumen for the cannula model.

After high-resolution data collection, standard helical refinement procedures led to a cryoEM reconstruction of the *ex vivo* cannulae at 2.3 Å global resolution (FSC 0.143 criterion; Figure 1a; Supplementary Figure 1f,g), which revealed two-strands having C2 symmetry, with a helical rise and twist along each strand of 3.18 Å and 12.1°, respectively, and a pitch of 94.6 Å. Given that the molecular identity of the *ex vivo* cannulae subunits was unknown, Modelangelo^41^ was used to build a *de novo* atomic model without providing an input protein sequence, followed by rounds of alternating manual and automated real-space refinement (see *Materials and Methods*). The final model revealed an intricate network of protein subunits that participated in polytypic radial and axial interactions (Figure 1b; Supplementary Figure 2). Specifically, the cannula protein subunits interlocked axially via N-terminal donor strand complementation (e.g. *i*→*j*→*k*; Figure 1b), and axially as well as radially via intermolecular calcium coordination (Figure 1c; Supplementary Figure 1h). Three types of calcium binding sites were identified that could be distinguished based on interfacial interactions between subunits. Type I and II binding sites mediated axial interactions between protomers (e.g., protomers *i* and *j*), and type III binding sites were located at the interface between two subunits that neighbored each other in the radial plane (e.g., the protomer interface between *j*-1 and *j*). Each calcium binding site type was unique and consisted of a combination of Ca^2+^ side-chain coordination via Asp or Glu residues, as well as main chain contacts with carbonyl groups and water molecules. We hypothesized that this elaborate network of calcium binding sites contributed to (i) the remarkable stability of the cannulae (*P. abyssi* is cultured at 100°C), but (ii) could also represent an environmental trigger for cannulae self-assembly (see below).

To identify the major subunit of the *ex vivo* cannulae fibers we first performed a hidden Markov search using the previously investigated cannula protein CanA (UniParc UPI0024181436)^35,42^ from *Pyrodictium abyssi* TAG11^32,33^ as query sequence against the *Pyrodictium abyssi* AV2 genome (NZ_AP028907.1). This search led to 19 putative CanA-like candidate proteins, 15 of which aligned onto a common consensus sequence (Supplementary Figure 3). Of those 15 sequences, 6 sequences were retained as plausible candidates based on sequence length arguments (i.e. primary mature sequence between 140 aa and 160 aa; Filter 1 in Supplementary Figure 4). The final assessment of the major subunit sequence was based on sidechain identification through manual inspection of the cryoEM map (Filters 2 and 3 in Supplementary Figure 4). WP_338251948.1 (annotated as “hypothetical protein”, InterProScan: no identified Pfam or PANTHER domains) was retained as the major subunit. Given the similarity (49.0% sequence identity) to previously identified CanA, and the fact that this sequence was derived from *ex vivo* analysis, we propose “CanX” as a naming convention. Interestingly, we found clear additional hexose-like density linked to residue Asn51 (Supplementary Figure 4). N-glycosylation in Thermoproteota was found to comprise a eukaryote-like GlcNAcβ(1-4)GlcNAc core structure.^43^ The excess density at Asn51 and the adjacent H-bond network fitted well with a β-linked N-acetylglucosamine (GlcNAc; albeit with some additional density of 3’), including additional density for a partially resolved hexose bound to its 4’ position (Supplementary Figure 4). We concluded that Asn51 is an N-linked glycosylation site, likely with GlcNAc core, which resulted in a band of glycans that helically wound around the cannula outer surface (Supplementary Figure 1i).

Intrigued by the number of detected CanA-like proteins in the *P. abyssi* genome, we inspected the genomic loci and found that 11 out 15 sequences were grouped into two extensive gene clusters, with the remaining 4 scattered across the genome (i.e. orphan sequences; Supplementary Figure 5). Interestingly, CanX was flanked by WP_338251946.1 (37.4% seq. id.) and WP_338251950.1 (29.45% seq.id.), which suggested that these three genes constituted an operon. Although we can only speculate on the function of the CanX homologues present in the genome at this point, we wondered if one or more of these CanX-like proteins could be (sporadically) integrated into cannulae (i.e., as minor subunits). The cryoEM volume shows no clear signs of subunit heterogeneity but if one or more CanX-like proteins were randomly integrated into the cannula lattice at low occupancy, then their impact on and contribution to the final 3D reconstruction would be limited. After inspecting the AlphaFold2^44^ structural predictions (Supplementary Figure 5), numerous candidates were found with substantial domain insertions, and we reasoned that such domains could act as fiducials to guide a particle classification process. To test that hypothesis, we tuned the filament tracer algorithm of cryoSPARC to selectively pick the edges of the cannula filaments and extracted the particles at a substantially smaller box size covering only 2.5 pitches to allow for greater sensitivity to local changes in downstream 2D classification jobs. Despite this approach, the resulting 2D class averages showed no signs of heterogeneity, nor could the presence of any additional domains be discerned as outward projections (Supplementary Figure 1j). We recognize that this approach cannot provide definitive proof that the cannula filaments were solely composed of CanX, but it suggested that if any other CanX homologues are incorporated into the cannulae, it would likely be at low sub-stoichiometric levels.

Next, we set out to determine the identity of the 2 nm fibrils that were associated with the cannula filaments (Figure 1d; Supplementary Figure 1b). Given the clear interaction between the two types of filaments, we referred to the 2 nm fibrils as putative cannula cargo. The reconstructed cryoEM volume showed no signs of the presence of the cargo, but that was expected due to the helical reconstruction method that was used, i.e., the cargo locally breaks helically symmetry and as a result any cargo density would be averaged out. To account for such a symmetry mismatch, we also performed C1 reconstructions followed by 3D classification but did not identify any 3D classes where the cargo is resolved, which was likely due to the absence of a specific cannula-cargo epitope (as already suggested by the raw micrographs). In the absence of a cryoEM resolved cannula-cargo interaction, we hypothesized that the cargo could be nucleic acid and collected a second cryoEM dataset after extended DNase I treatment. The resulting raw micrographs showed no signs of the presence of any cargo after DNase I treatment. To formalize that observation, we adopted the following workflow. Starting from high resolution helical reconstructions of both DNase I treated and non-treated particles, we used the respective consensus volumes as input in particle subtraction jobs – the rationale being that particle alignment and classification would be dominated by the cannula signal for the non-subtracted particles. Next, the subtracted particles were 2D classified and inspected for the presence of fibrillar classes that would match the dimensions of the cargo in the raw micrographs. For the non-treated cannulae dataset, such cargo 2D class averages were produced as evidenced from the apparent diameter (i.e. 2 nm) and the mapping of the particles-corresponding to the cargo 2D class averages-back onto the raw micrographs, which confirmed that these corresponded to boxed regions with cargo clearly present. In contrast, no cargo 2D class averages could be obtained following DNase I treatment, which allowed us to conclude that the cargo fibrils were DNA. Although the cryoEM images were suggestive of a luminal location, these did not unambiguously exclude the possibility that the cargo to be bound to the cannulae surface. However, indirect evidence further suggested a luminal interaction based on the surface electrostatics of the cannula filaments. Cannulae displayed a highly polar structure that has a predominantly negative outer surface and a highly positive lumen (Figure 1d). The presence of a positively charged lumen for a negative cargo might seem contradictory from the perspective of DNA transport but could be rationalized through a mechanism of (i) counterion mediated charge neutralization or (ii) the involvement of effector molecules bound to the DNA to facilitate transport through the cannulae fibers.^17–19,45^ Taken together, these data suggested that cannulae might function as molecular conduits between cells for the exchange of DNA.

In addition to the cannulae fibers, a second class of pili was also found to be present (Supplementary Figure 1d). These pili existed as large super bundles with a local undulating character that was reminiscent of the structure of the archaeal bundling pili (ABP) of *Pyrobaculum calidifontis* (PDB: 7UEG).^10^ Although the 2D class averages of bundling pili don’t show homogenous packing of the filaments, the resulting power spectrum showed a pronounced maximum at 1/47Å, which we presumed corresponded to the axial rise of the subunits in the individual pili. For ABP, however, that axial rise is 33 Å and the major subunit is AbpA (WP_011849593.1). Using HMMer^46^ and Foldseek,^47^ we found no putative homologues of AbpA in the genome of *Pyrodictium abyssi* AV2. AbpA was distantly related to TasA (UniProt: P54507), the major biofilm matrix component of Gram-positive *Bacillus subtilis*, which was shown to have an axial rise of 48 Å (PDB: 8AUR).^9^ Using TasA as a query, we identified the hypothetical protein WP_338249486.1 (AAA988_RS09245) in a structural similarity search (Supplementary Figure 7). Although the sequence identity to TasA is low (16%), the AlphaFold3^48^ prediction of a WP_338249486.1 dimer revealed notable structural similarities, i.e. WP_338249486.1 is predicted to form a donor strand complemented dimer with a monomer-to-monomer distance of 47 Å, which agreed with the experimentally determined axial rise of the bundling pili of *P. abyssi* AV2 (Supplementary Figure 1d,e). Looking at the local genomic context to AAA988_RS09245, we identified AAA988_RS09250, presumed to form an operon with AAA988_RS09245, and predicted to encode for a signal peptidase I. Based on these data we hypothesized that the bundling pili found in the extracellular milieu of *P. abyssi* are formed by

WP_338249486.1, and we propose the following nomenclature, AbpX. While never previously identified in EM analyses of *Pyrodictium* extracellular filaments, the AbpX fibers were more similar in structure to the taxonomically more distant TasA filaments than to previously characterized AbpA pili from *P. calidifontis*. It is remarkable that two different precursors proteins, CanX and AbpX, polymerized into structurally distinct extracellular filaments through a common process of donor strand complementation on the surface of *P. abyssi*.

While the cryoEM structural analysis of the CanX filament provided insight into the remarkable stability of *P. abyssi* cannulae, the challenging culture conditions would preclude large-scale preparations of cannulae for biomaterials applications. Previous experimental investigations demonstrated that recombinant expression of cannula-like proteins in heterologous bacterial hosts represented a viable preparative route to soluble precursors of structurally similar filamentous assemblies.^35,42^ CanA, a cannula-like protein from *P. abyssi* strain TAG11, has been the most thoroughly studied of these recombinant precursors.^32^ Previous investigators reported that recombinant CanA could assemble *in vitro* into tubular filaments in the presence of calcium ions, in which the resultant filaments displayed a similar diameter and morphology to *ex vivo* cannulae from *P. abyssi* TAG11 under low resolution electron microscopy imaging.^35^ In addition, a recombinantly expressed truncation of CanA, K_1_-CanA, corresponding to a deletion of the first ten amino acid residues of the donor strand of the mature protein, displayed no polymerization propensity under similar conditions.^42^ The induction of self-assembly of recombinant CanA in the presence of calcium ion and its inhibition upon partial deletion of the N-terminal donor strand suggested that recombinant cannula-mimetic proteins could polymerize *in vitro* through a mechanism that mimicked the assembly of the *ex vivo* filaments.

A structural determination of *in vitro* assemblies of recombinant CanA was undertaken to compare to the corresponding structure of the *ex vivo* filament. A codon-optimized version of the coding sequence of the mature CanA protein, i.e., after deletion of N-terminal twenty-five amino acids of the signal peptide sequence, was designed for bacterial cell expression (Supplementary Figures 8a and 9a). Previous research demonstrated that this CanA protein variant could be expressed and purified as a soluble protein in *E. coli* host systems in good yield (2.1 g protein/250 g wet cell weight).^35^ Following the published procedure, recombinant CanA was obtained in an unoptimized yield of 31 mg of purified protein per L of culture from the cell lysate after expression in *E. coli* strain BL21Gold(DE3) (see *Materials and Methods*). The purity and identity of the recombinant CanA protein were confirmed using SDS-PAGE analysis and electro-spray ionization (ESI) mass spectrometry (Supplementary Figures 10a and 11a). Sedimentation velocity analytical ultracentrifugation indicated that CanA migrated as a monomer under these conditions at an apparent molar mass that agreed with mass spectrometric analysis and the monoisotopic mass calculated from the amino acid sequence of CanA (Supplementary Figure 12a). Thermolysis (50° - 80°C) of solutions of purified CanA in a low-salt buffer (50 mM Tris-HCl, pH 7.5, 80 mM NaCl, 0.1 mM, EDTA, 9% glycerol) led to rapid polymerization in the presence of CaCl_2_ (20 mM).^35^ Negative-stain TEM images of these solutions revealed the presence of a structurally uniform population of high aspect-ratio tubular assemblies of ∼25 nm in diameter (Supplementary Figure 13a), which agreed with previous reports from EM imaging of *ex vivo P. abyssi* TAG11 cannulae and recombinant polymers of CanA.^32,35,38^ The cannula-like filaments of CanA laterally associated into large rafts like those observed in preparations of *ex vivo P. abyssi* AV2 cannulae (see above). Similar behavior was observed for cannulae under native growth conditions in cultures of *P. abyssi* TAG11, in which fast freeze/deep etch electron microscopy revealed the presence of bundles containing up to a hundred cannulae at the cell surface.^32^

CryoEM analysis was employed to determine the structure of the recombinant CanA assemblies. Despite extensive lateral association between filaments observed in the raw micrographs, a sufficient number of isolated particles could be classified as the basis for helical reconstruction (Figure 2a). The 2D class averages confirmed that the tubular filaments were uniform in inner and outer diameter, ∼18 nm and ∼27 nm, respectively, despite *in vitro* assembly conditions that differed significantly from the native growth environment of *P. abyssi*. A 3D volume was reconstructed from the 2D projection images using iterative helical real-space reconstruction (IHRSR)^49,50^ after determination of helical symmetry (Supplementary Figure 14a).^25^ An unambiguous atomic model could be built into the 2.6 Å resolution 3D density map (Figure 2b; Supplementary Figure 14b, c). In contrast to the C2 symmetry in the *ex vivo* cannulae, the CanA filament was a C1 assembly of protomers along a left-handed 1-start helix with a rise of 1.59 Å and a twist of-173.75°. The most prominent structural feature of the CanA filament, as for the CanX filament, was the presence of right-handed 2-start helices that generate a ∼4.6-4.7 nm periodicity (Figures 1a and 2b,c), i.e., half the pitch of the 2-start helices, which was consistent with previously reported TEM images of native cannulae and recombinant cannula-like tubes.^32,35,38^

**Figure 2:**
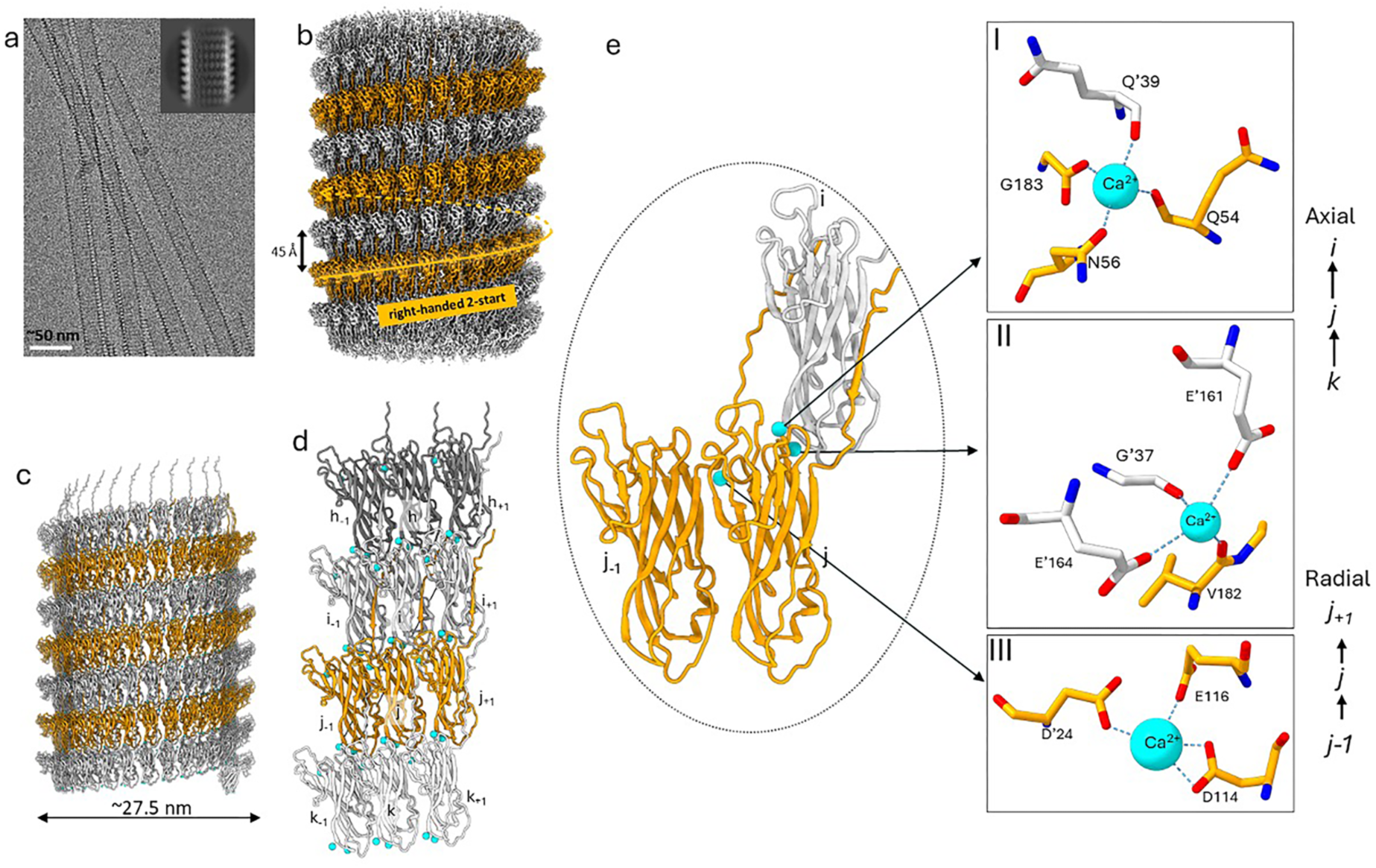
CryoEM structure of *in vitro* cannula-like tubules formed by the self-assembly of recombinant CanA. (a) representative raw cryoEM micrograph and 2D-class average of the recombinant CanA filament; (b) cryoEM volume depicting right-handed 2-start helical packing under C1 symmetry (rise and twist of 1.59 Å and-173.75°, respectively); (c) helical model of CanA filament; (d) local interaction network between neighboring CanA subunits with three Ca^2+^ ions bridging protomers in a 12mer segment of the cannulae; (e) close-up view of three Ca^2+^ ions binding sites in stick representation.

As observed for the *ex vivo* filament, the atomic model of the CanA tubules suggested that donor strand complementation was the primary mechanism that drove polymerization of CanA into cannula-like filaments and stabilized the resultant assemblies. The relative orientation of protomers in the polymer, e.g., along the *h→i*→*j*→*k* direction (Figure 2d) was defined by a rotation of 1.25° and an axial rise of 46 Å (1.5° and 48 Å, respectively, for the corresponding interaction in CanX). Successive protomers were aligned nearly parallel to the central axis of the helical assembly (Figures 1b and 2d). Donor-strand complementation linked protomers located on structurally adjacent 2-start helices (Figure 2b, c), and reinforced the structural interfaces within the CanA filament. The cannula-like structure of CanA displayed an extensive network of inter-protomer contacts that involved subunits located as far away as *j±60* on either side of a central protomer (Supplementary Figure 14d). PISA (Protein, Interfaces, Surfaces, and Assemblies) analysis^51^ indicated that, while donor strand complementation between protomers buried the greatest amount of surface area, significant interactions also occurred between subunits located within the same protofilament (e.g., subunits *j-1* and *j*) along a 2-start helix (Figure 2d). In addition, as observed in the *ex vivo* filament, three calcium ions could be fit into the EM density map of the *in vitro* filament (Figure 2d). The calcium ion binding sites were structurally homologous to those of the *ex vivo* filament, mediating similar polytypic radial and axial interactions between structurally adjacent protomers (Figure 2e). Other than the observed helical symmetry, the most significant difference between the *ex vivo* and *in vitro* cannula filaments was the structure of the protomers. While the core structures of CanX and CanA were based on homologous jelly-roll folds (49.0% seq. ID), the presence of extended loops were observed in the CanA protomer, and its corresponding AlphaFold3^48^ predicted structure, which were not present in CanX (Supplementary Figure 15a). In the recombinant CanA structure, these loops projected from the outer surface of the filament, which resulted in a pair of ridges that coincided with the 2-start helices (Figure 2b, c). While the CanX protomer lacked these loop insertions, a cannula-like protein (WP_338251966.1) was identified at a different locus in the *P. abyssi* AV2 genome (Supplementary Figure 5) in which the predicted structure was homologous to CanA (51.1% seq. id.). However, outer surface projections that could account for the presence of this CanA-like protein were not detected in the *ex vivo* filament (see above).

The conservation of the calcium binding sites in the structures of the *ex vivo* and *in vitro* cannula filaments, as well as its induction of CanA assembly, raised a question regarding relationship between calcium ion binding and donor strand complementation. The polymerization of the cannula-like proteins resulted from progressive donor strand complementation in concert with calcium ion binding. However, for CanA, calcium ion binding was a necessary but not sufficient criterion for polymerization. While the monomer was competent for polymerization in the presence of calcium ions, the process was kinetically slow at ambient temperature and required thermolysis to promote assembly. Under native growth conditions for *P. abyssi* (∼100°C), the temperature would be sufficient to initiate polymerization of cannula-like proteins since the calcium concentration in seawater (400 ppm, ∼10 mM) would be within the permissible concentration range. Hypothetically, other divalent metals ions present in seawater could promote self-assembly assembly of cannula-like proteins into filaments at the cellular surface of *P. abyssi*. The Mg^2+^ ion, the most abundant divalent cation in seawater (1,300 ppm, ∼54 mM), has a smaller ionic radius than Ca^2+^ (72 pm versus 100 pm) and was not able to induce filamentation of recombinant CanA *in vitro* under equivalent conditions (20 mM ion, 50-80°C, low salt buffer, pH 7.5). Notably, the addition of terbium ion (10 mM), which is often employed as a calcium ion surrogate in biophysical studies of calcium binding proteins (Tb^3+^ ionic radius ∼106 pm),^52^ rapidly initiated polymerization of recombinant CanA into cannula-like filaments at ambient temperature (*VPC, unpublished data*). EDTA (10 mM), a strong chelator of calcium ion (log*K_f_* ∼10.65), inhibited the Ca^2+^ induced polymerization of CanA into cannula-like filaments. However, treatment of pre-formed cannulae with EDTA did not induce disassembly unless the solution was heated to 60°C. SV-AUC measurements indicated that CanA remained a monomer indefinitely in solution in in the presence of sub-critical concentrations of Ca^2+^ ion below the optimal polymerization temperature range (Supplementary Figure 10a)

To gain insight into the potential role of calcium ion in driving filament formation, single-crystal X-ray diffraction was employed to determine the structure of the monomeric form of CanA in the absence of calcium ion (Figure 3a). A comparison between the crystal structure of the CanA monomer and the corresponding CanA protomer in the assembly revealed significant conformational differences between the two structures that were localized to protein segments associated with subunit-subunit interfaces in the filament. The primary difference lay in the conformation of the *N*-terminal nineteen amino acids in CanA. These residues were not observed in the crystal structure of the CanA monomer, which suggested that the *N*-terminal segment was disordered and did not contribute to the diffraction. In contrast, these *N*-terminal amino acids adopted a β-strand conformation (strand A) in the cryoEM structure of the CanA protomer, which mediated the donor strand-acceptor groove interaction between axially adjacent protomers (e.g., *i,j*) in the filament (Figure 2b, c). In addition, significant structural differences were observed in the conformation of two loops, corresponding to residues 35-39 and 161-173, that comprised part of the axial Ca^2+^ binding sites (Figure 3b) within the cannula filament. In the crystal structure, the electron density corresponding to residues 35-39 was absent, which implied that the loop conformation was dynamic in the absence of calcium ion. In addition, a backbone alignment of the CanA crystal structure and the CanA cryoEM protomer structure indicated that a loop region corresponding to residues 161-173 was significantly displaced between the monomeric and assembled forms of CanA (Figure 3c). We attributed the observed conformational changes in these two loops to the absence or presence of calcium ion binding. Alignment of the monomeric CanA structure to an axially interacting pair of protomers (*i,j*) suggested a mechanism whereby calcium ion binding could promote donor strand complementation (Figure 3d). In the structure of the CanA filament (Figure 3e), calcium ion binding stabilized the conformation of the two loop regions and promoted a local interaction between them that exposed the acceptor groove of the monomer. In the crystal structure of the monomer, loop region 161-173 adopted a conformation that occluded the acceptor groove, which prevented it from accepting the donor strand of another monomer and precluded donor strand complementation potentially preventing filament formation (Figure 3f). We propose that calcium ion binding to the monomer triggered a structural rearrangement that rendered the monomer competent for subsequent donor-strand complementation and polymerization into a cannula-like filament. However, the rate at which polymerization occurred was observed to depend on temperature, perhaps mediated through a thermally activated process that was not apparent from the available biophysical data. The crystal structure of the monomer clearly shows that the N-terminal region is free and available to invade another subunit when polymerized, and not self-complemented as part of a β-sheet.

**Figure 3:**
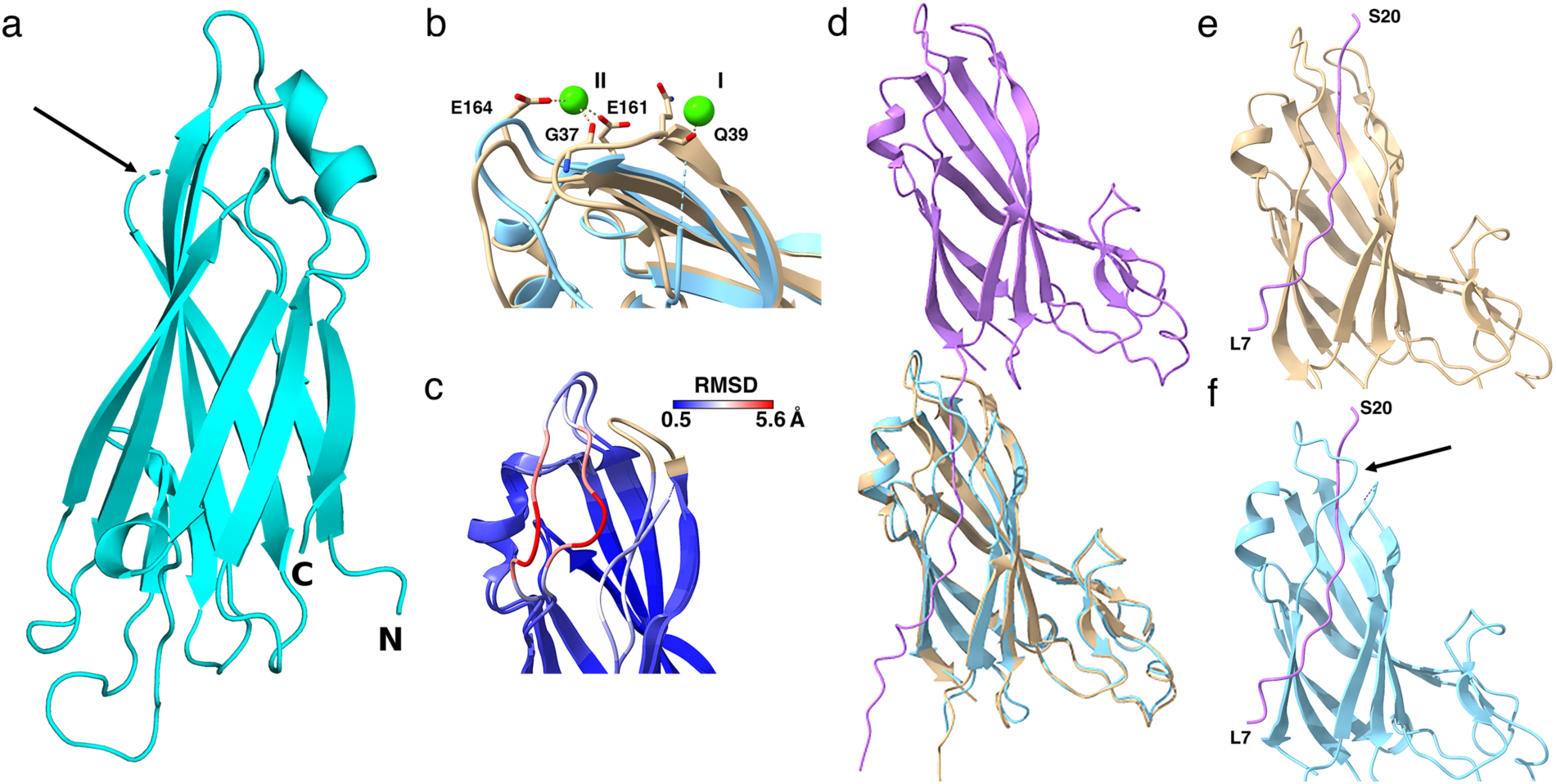
Crystal structure analysis of the CanA monomer. (a) crystal structure of the CanA monomer (cyan) depicted as a ribbon diagram. The arrow indicates a rendering of the disordered loop region 34-40 (b) close-up of the superposed CanA monomer from the crystal structure (blue) with the cryoEM CanA protomer (tan) depicting the significant conformational change within loop region 161-173. (c) close-up of the RMSD rendering from a Matchmaker backbone alignment between the CanA protomer and monomer that highlights the difference in loop region 161-173. (d) alignment of the CanA monomer from the crystal structure (blue) to an interacting axial pair of CanA protomers from the cryoEM structure. The N-terminal donor strand from protomer *i* (violet) inserts into the acceptor groove of protomer *j* (tan). (e)-(f) zoomed-in views of loop region 161-173 in the CanA cryoEM structure (e) and the superposed CanA monomer (f). The latter view revealed a potential steric clash (arrow) between this region of the CanA crystal structure with donor strand residues Tyr11 to Ser20 that could conceivably preclude CanA oligomerization in the absence of calcium ion binding.

To identify other potential cannula-mimetic proteins in related archaeal species, Foldseek was employed with the protomer structure of CanA (PDB: 7UII) serving as a query.^47^ Several sequences were identified within the family *Pyrodictiaceae*, including five proteins from the related species *Pyrodictium occultum* and four putative proteins derived from metagenome-assembled genomic (MAG) data reported for an uncultured *Hyperthermus* species (Supplementary Table 1). AlphaFold2 structural predictions were available for the respective proteins through the UniProt webserver (www.uniprot.org).^53^ In each case, the presence of a jelly-roll fold was predicted for the respective monomeric structures with high pLDDT and pTM, although poorly predicted insertions were also observed for several of the larger proteins as observed for *P. abyssi* (Supplementary Figure 5). In most cases, an unstructured N-terminal domain was predicted for the structures of the mature protein sequences. As determined for CanA and CanX, most of these cannula-mimetic protein sequences contained potential N-glycosylation sequons (NxS/T) that localized to surface loops in the AlphaFold predictions.

Two proteins from *Hyperthermus* were chosen for investigation since they were derived from a different genus in the family *Pyrodictiaceae*. These proteins, Hyper1 (hypothetical protein DSY37_00495, UniProt: A0A432RA20) and Hyper2 (hypothetical protein DSY37_03180, UniProt: A0A432R7L7), shared a pairwise sequence identity of approximately 37.8% with each other and approximately 27.2% and 23.4% with CanA, respectively. Codon-optimized coding sequences (Supplementary 9b) of the corresponding proteins were cloned into the plasmid pD451-SR and transformed into *E. coli* strain BL21Gold(DE3). Expression cultures were screened for the presence of the respective proteins under IPTG induction using the procedure optimized for recombinant CanA (*see Materials and Methods*). While SDS-PAGE analysis indicated that both proteins were successfully expressed, the Hyper1 protein accumulated in inclusion bodies that could not be refolded after denaturation and was not further studied. The Hyper2 protein accumulated in the soluble fraction of the cell lysate and could be purified at an isolated yield of 80-100 mg protein/L of cell culture using the procedure developed for CanA. The purity and identity of the recombinant Hyper2 protein were confirmed using SDS-PAGE analysis and electro-spray ionization (ESI) mass spectrometry (Supplementary Figures 10b and 11b). As observed for CanA, sedimentation velocity AUC analysis indicated that Hyper2 behaved as a monomer in the absence of calcium ion (Supplementary Figure 12b).

In contrast to CanA, Hyper2 assembled rapidly at ambient temperature in the presence of calcium ion (≥ 10 mM) without thermolysis. However, negative-stain TEM analysis (Supplementary Figure 13b) indicated a range of filamentous assemblies including single-walled tubular filaments of different diameter and multiple-walled tubes. The structural polymorphism of the Hyper2 assemblies was confirmed in the raw cryoEM micrographs (Figure 4a, Supplementary Figure 16a). Helical reconstruction was employed to structurally analyze two of the single-walled tubes corresponding to diameters of 240 Å and 270 Å after independent 2D classification and assignment of helical symmetry (Supplementary Figures 16b). Standard helical refinement procedures led to cryoEM reconstructions (Figure 4b and Supplementary Figure 16c) at a 3.5 Å global resolution for both classes of filaments (FSC 0.143 criterion; Supplementary Figure 17b, c, Supplementary Table 2). Reliable atomic models could be built into the respective EM density maps for the two classes of Hyper2 filaments, which revealed that they were structurally related but based on different helical symmetries. The structure of the thick tube displayed C8 symmetry with a helical rise of 11.16 Å and twist of 9.59°, while the structure of the thin tube revealed C1 symmetry with a helical rise of 1.42 Å and a twist of 46.23° (Figure 4b, c and Supplementary Figure 16c). The most prominent structural feature of the two Hyper2 assemblies was the presence of eight protofilaments. As observed for the *Pyrodictium* cannulae reconstructions, the Hyper2 filaments were linked through a combination of donor strand polymerization between protofilaments and calcium ion binding interactions (Figure 4d, e; Supplementary Figure 16d). The distance between the protofilaments along the direction of donor strand polymerization was ∼4.6-4.7 nm for the Hyper2 cannulae, which compared well to that observed for the CanX (∼4.8 nm) and CanA (∼4.6 nm) filaments. The most significant structural difference observed between the *Pyrodictium* and *Hyperthermu*s cannulae filaments lay in the number and geometrical arrangement of the protofilaments. For the *Hyperthermus* cannulae, eight protofilaments propagated along right-handed 8-start helices having a periodicity of ∼5.2 nm (one-eighth of the 8-start helical pitch) and twists of 9.59° (C8) and 9.84° (C1), respectively (Figure 4b and Supplementary Figure 16c). While for the *Pyrodictium* cannulae, two protofilaments were aligned along right-handed 2-start helices having a periodicity of 4.6-4.7 nm (Figures 1a and 2b, c). In all cases, donor strand complementation occurred between structurally adjacent protofilaments and reinforced the supramolecular structure of the respective filaments. The donor strand-acceptor groove interaction occurred along a series of left-handed 31-start (C1) or 32-start (C8) helices for *Hyperthermus* cannula-like filaments (Figure 4d; Supplementary Figure 16c). In contrast, the donor strand polymerization in the *Pyrodictium* cannulae occurred along a set of right-handed 30-start helices for CanX (Figure 1c) or 29-start helices for CanA (Figure 2d). The propagation axis of the donor strand polymerization was nearly parallel to the central helical axis for the *Pyrodictium* cannulae (∼4° tilt), whereas donor strand propagation was tilted at an angle (∼18°) from the central axis in the Hyper2 assemblies.

**Figure 4:**
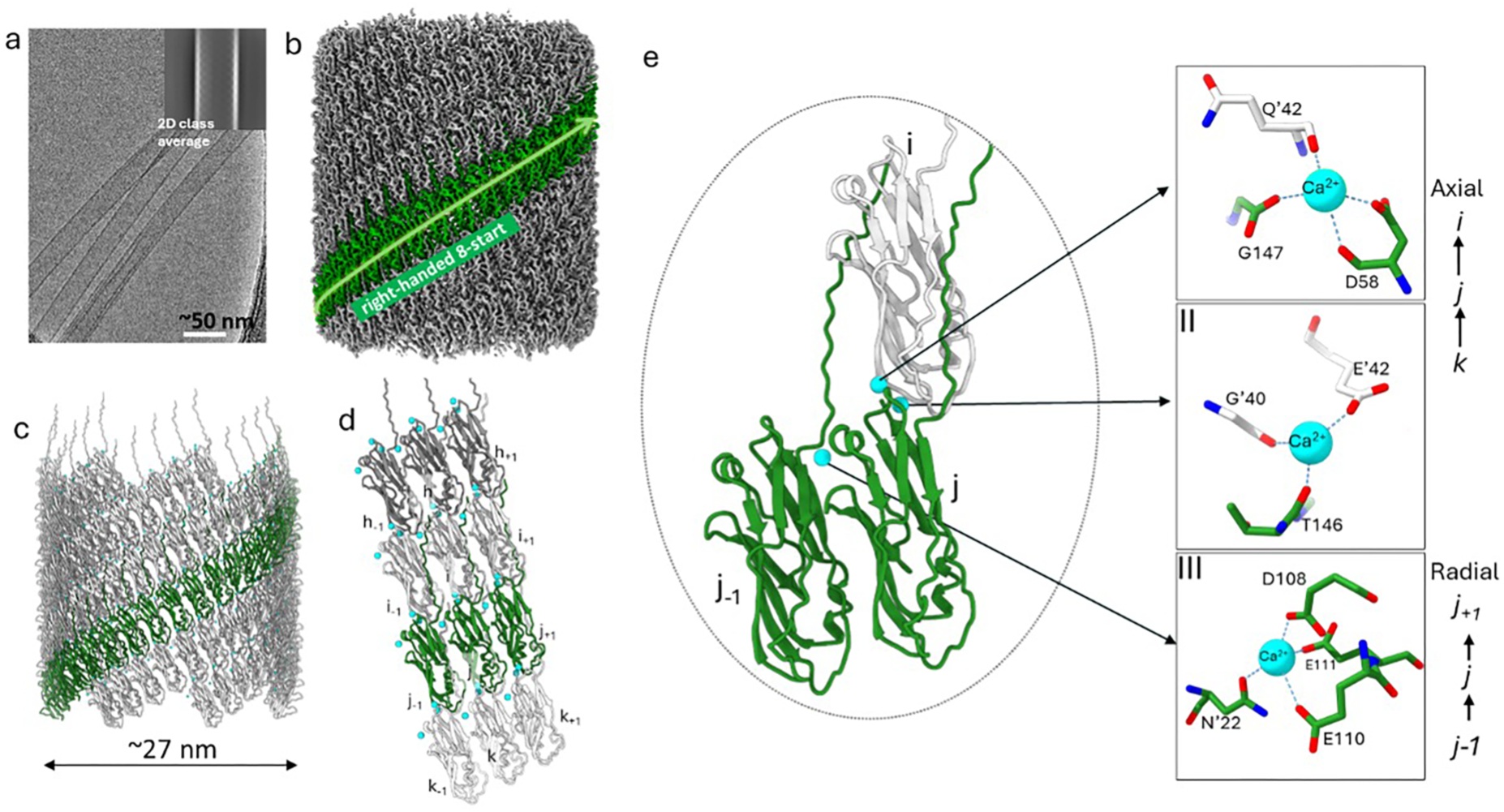
CryoEM structure of cannula-like tubules (Hyper2 thick tube) formed by the self-assembly of recombinant Hyper2. (a) representative raw cryoEM micrograph and 2D-class average of recombinant Hyper2 tubes; (b) cryoEM map showing right-handed 8-start helical packing with C8 rotational symmetry (rise and twist of 11.16 Å and 9.59°, respectively); (c) helical model of Hyper2 thick tube; (d) local interaction network between neighboring Hyper2 subunits with three Ca^2+^ ions bridging protomers in a 12mer segment of the cannulae; (e) close-up view of the three Ca^2+^ ions binding sites in stick representation.

Despite the differences in helical symmetry, the protomer structure and local structural interfaces were remarkably conserved between the *Hyperthermus* and *Pyrodictium* cannulae. The core structure of the protomer was based on a donor-strand complemented jelly-roll fold in which calcium ions were observed to bind at the radial and axial interfaces between protomers. Structural alignment of the Cα backbone atoms of the CanX, CanA, and Hyper2 protomers suggested a close correspondence between the respective jelly-roll folds despite the presence of two large loops in the CanA structure (Supplementary Figure 15a). Two distinct dimeric structural interfaces were identified in the local interaction network of the respective filaments (Supplementary Figure 2), which were associated with radial interactions between laterally adjacent protomers within the same protofilament (e.g., subunits *j-1,j,j+1*) and axial interactions between protomers located in different protofilaments (e.g., subunits *i,j,k*). Pairwise backbone alignments of the dimeric radial and axial interfaces between protomers in the different cannulae revealed that the structures were strongly conserved (RMSD ≈ 0.8-1.2 Å, Figure 15b). This strong conservation of interfacial interactions among the cannulae can be contrasted to the limited sequence identity between the *Pyrodictium* and *Hyperthermus* cannula-like proteins (Figure 15c).

As in the *ex vivo* and *in vitro P. abyssi* cannulae, three calcium ion binding sites were observed per protomer in the Hyper2 filaments at structurally homologous positions within the assemblies. A multiple sequence alignment (MSA) of the mature CanX, CanA, and Hyper2 proteins indicated that the residues in the calcium ion binding sites were conserved (Figure 5a), which suggested a similar role for calcium ion in inducing assembly and maintaining the structural integrity of the respective cannulae (Figures 1b,c; 2d,e, and 4d,e). Donor strand complementation served as the primary mechanism through which cannula-like protein monomers underwent polymerization into cannula-like filaments. Donor strand A of subunit *j* forms a series of hydrogen-bonded interactions with strands B and K of the acceptor groove on an axially adjacent subunit *i* (shown for CanX filament in Figure 5b). The hydrophobic groove of strands B and K define a series of five pockets, P1-P5, which accommodated residues on one face of the donor strand (Figure 5c-e). The donor strands of the cannula-like proteins displayed an amphipathic sequence pattern in which hydrophobic residues alternated with hydrophilic residues. The sidechains of the former residues were buried in the acceptor groove while the polar sidechains of the latter residues were exposed on the filament surface. The donor strands of the cannula-like proteins displayed significant sequence homology, in which an aromatic residue (F/Y) occurred at the site that occupied P1, while smaller sidechain hydrophobic residues (G/A/V) occupied the sites on strand A that interacted with P2-P5 (Figure 5c-e). The central Gly residue (cyan, Figure 5c-e) was conserved between all identified cannula-like proteins in *Pyrodictium* and *Hyperthermus* species. As observed for classical chaperone-usher (CU) pili,^3,4,8,54,55^ the presence of a conserved glycine residues might be present to ensure the correct register of the hydrophobic knobs in holes interaction of donor strand-acceptor groove interaction between cannula protomers. To test the importance of the glycine residue’s role in donor strand complementation, a G13A mutant of the Hyper2 protein was prepared. The G13A mutant was severely compromised in its ability to form filaments in the presence of calcium ion when compared to the *wild-type* Hyper2 protein under identical conditions.

**Figure 5:**
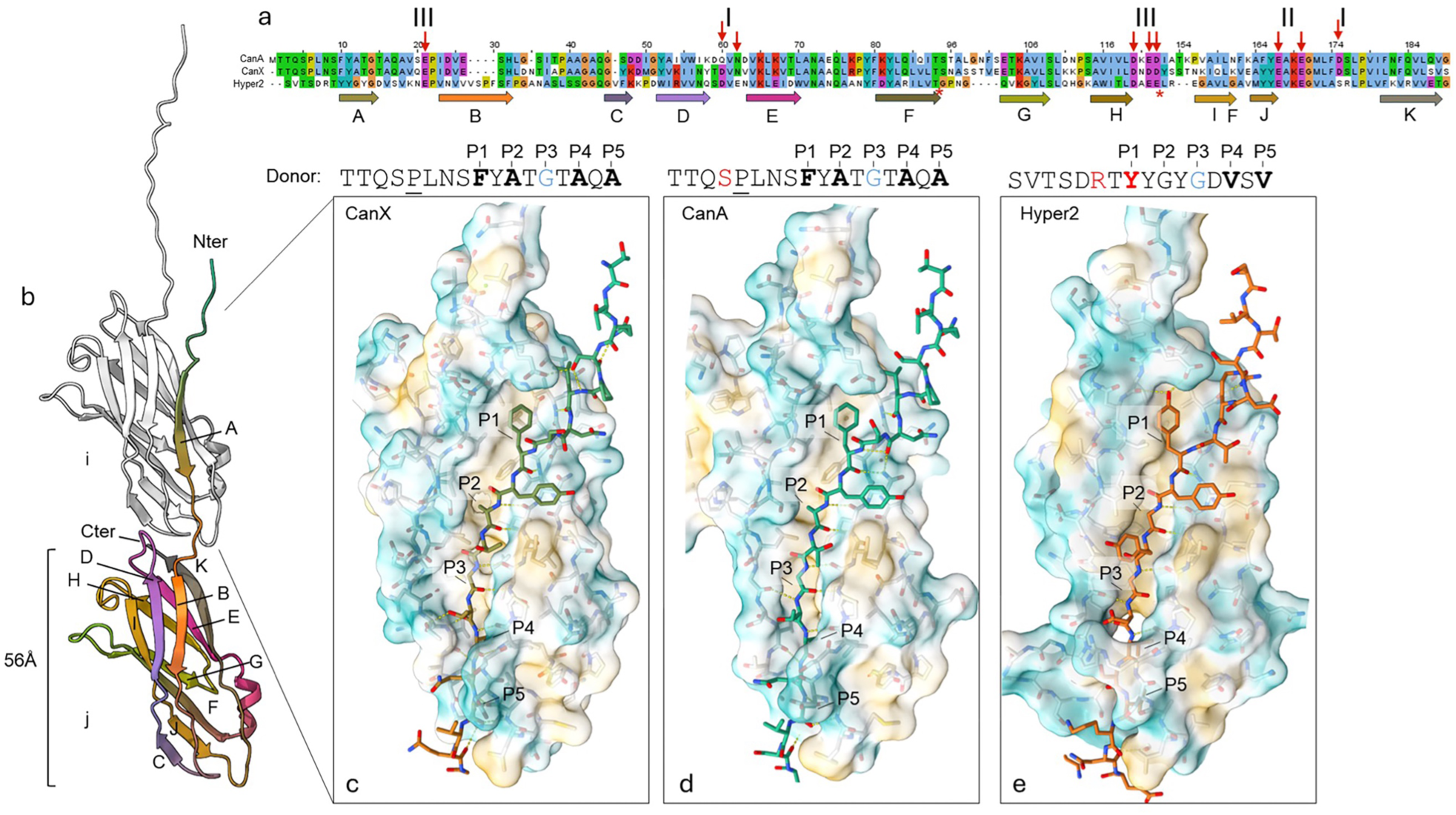
**Conserved donor strand complementation across cannulae forming subunits**. (a) multiple sequence alignment (MSA) of mature CanA, CanX and Hyper2 sequences (Jalview, Clustal coloring). Red stars denote regions where CanA loop/domain insertions were removed for visual clarity. Red arrows highlight conserved residues that interact with calcium in type I, II and III coordination centers. Arrows below the MSA denote secondary structure elements color coded according to the monomer cartoon representation in (b); (b) Illustration of the cannula donor strand complementation mechanism shown here for CanX: the N-terminal strand A of subunit j docks into the surface exposed hydrophobic groove defined by the region between strands B and K of subunit I; (c)-(e) zoomed-in renditions of the donor strand (stick representation) docking into the respective B/K grooves for CanX, CanA and Hyper2, respectively. Donor sequences above panels (c)-(e): hydrophobic residues docked into complementary pockets highlighted in bold; residues forming sidechain-mediated intermolecular hydrogen bonds shown in red; proline directing the N-terminus towards the ‘lattice contact’ underlined.

This structural similarity between cannulae and the mechanistically unrelated chaperone-usher pili prompted us to systematically evaluate the evolutionary relationships of CanX and AbpX with other bacterial and archaeal cell surface filaments formed through donor strand complementation.^3–7,9–11,39^ Based on HMMer searches and signal peptide prediction, all CanX homologs are exclusively found in the *Pyrodictiaceae* family and possess a Sec/SPI signal sequence, which suggested they are transported by the Sec translocon and subsequently cleaved by Signal Peptidase I, like other characterized donor strand complemented filaments. However, the genomic neighborhood of the genes encoding CanX homologs lacked genes encoding putative chaperones or dedicated signal peptidases, supporting the notion that CanX could self-assemble in the presence of calcium without external assistance. Similarly to CanX, AbpX also possessed a Sec/SPI secretion signal peptide. However, HMMer searches revealed that AbpX homologs were more widely distributed across archaea,^10^ including in *Pyrodictiaceae* species that lacked CanX homologs, such as *P. delaneyi* Su06^34^ (WP_055409108.1). The searches with AbpX and its archaeal homologs also identified sequence matches to bacterial TasA family proteins, with pairwise sequence identity levels of 30%-35%, which supported the classification of AbpX within the TasA superfamily (Figure 6a, b). Like bacterial TasA proteins, AbpX homologs in archaea were frequently found adjacent to genes encoding signal peptidase I or other AbpA-/TasA-like proteins, underscoring the evolutionary and functional connections between these systems.

**Figure 6:**
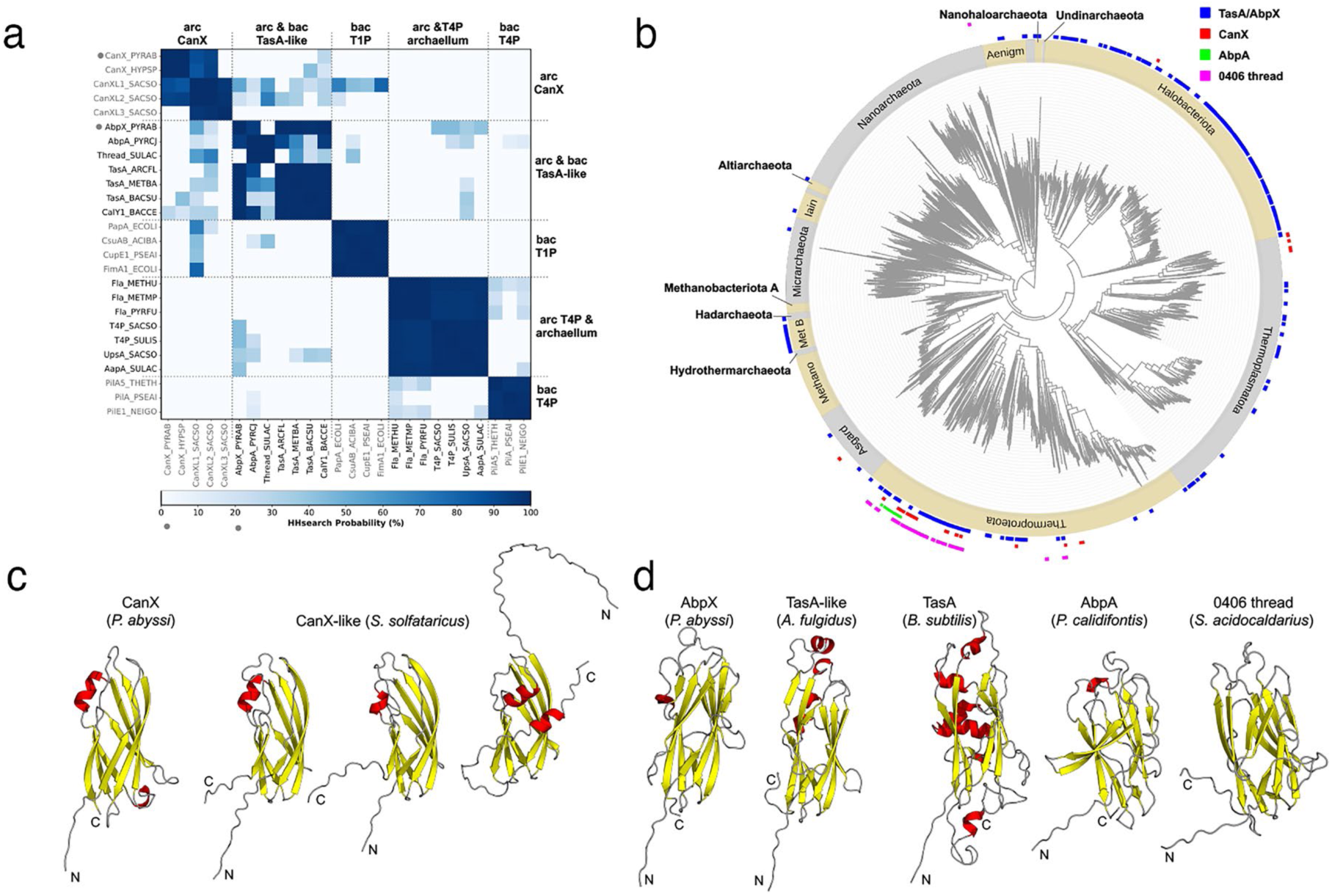
**CanX and AbpX belong to the TasA superfamily**. (a) pairwise profile HMM comparison of representative archaeal and bacterial cell-surface filament subunits. The heatmap shows HHsearch probabilities (%) for each pairwise comparison. The NCBI/UniProt accession numbers of the proteins are provided in the Methods section. (b) distribution of the TasA superfamily across archaea, visualized on a species-level archaeal tree from the Genome Taxonomy Database. Colored dots indicate the presence of specific protein families: TasA/AbpX (blue), CanX (red), AbpA (green), and *S. acidocaldarius* 0406 thread (magenta). Phyla are indicated on the outer ring of the tree. (c) and (d) structural representations of representative TasA superfamily proteins. AlphaFold3 models are shown for selected proteins, while cryoEM structures are displayed for *P. calidifontis* AbpA (PDB: 7UEG) and the *S. acidocaldarius* 0406 thread subunit (PDB: 7PNB). α-helices are colored red, and β-strands are colored yellow.

To uncover potentially distant CanX homologs that HMMer searches might have missed, sensitive profile HMM-based sequence searches were employed using HHpred. We searched the profile HMM databases of proteomes from several representative archaea for homologs of CanX, which identified three distant homologs in *Saccharolobus solfataricus* (WP_009990924.1, WP_009991716.1, WP_029552507.1) with HHpred probability values of greater than 90% (Supplementary Figure 18a). Despite sharing only ∼18% pairwise sequence identity with *P. abyssi* CanX, the structures of these proteins, predicted by AlphaFold2, retained the jelly-roll fold, showing a striking structural resemblance to CanX (Figure 6c). Homo-oligomers of these proteins, as predicted by AlphaFold2-multimer,^56^ suggested their potential to polymerize through donor-strand complementation (Supplementary Figure 18b). These proteins were employed for subsequent HMMer searches, which identified over 100 CanX-like proteins in archaea, mostly within the phylum *Thermoproteota*, specifically in the *Sulfolobaceae* and *Thermocladiaceae* families (Figure 6b). For instance, we identified a homolog each in *Vulcanisaeta distributa* DSM 14429 (WP_054853146.1) and *Caldivirga maquilingensis* IC 167 (WP_156769888.1), two in *Sulfolobus acidocaldarius* DG1 (WP_011279025.1, WP_011279026.1), and four in *Metallosphaera yellowstonensis* MK1 (WP_009071828.1, WP_009072785.1, WP_009072754.1, WP_009071324.1). AlphaFold3 models for these homologs suggested that they could also adopt the jelly-roll fold (Supplementary Figure 18b) and could undergo donor strand complementation.

The jelly-roll fold of cannula-like proteins was a structural feature that was shared with the bacterial TasA protein and its recently identified archaeal homolog, AbpA, from *P. calidifontis*. To explore the evolutionary relationships among these proteins, HHpred was employed for pairwise sequence comparisons and searches against the PDB and Pfam profile HMM databases. These searches identified statistically significant matches (HHpred probability values between 80%-95%) between CanX-like proteins and AbpA (PDB: 7UEG), the *S. acidocaldarius* 0406 filament subunit (PDB: 7PNB), as well as the Pfam Peptidase_M73 family (PF12389), which encompassed bacterial TasA homologs. The pairwise sequence identity across these matches was low (below 20%), indicating that CanX and CanX-like proteins form a highly divergent family within the TasA superfamily. The *S. acidocaldarius* 0406 thread subunit, which was previously proposed as a representative of a novel class of archaeal filaments, also showed statistically significant matches to CanX-like proteins, AbpA, and TasA family proteins in our HHpred searches. However, the low sequence identity among these proteins (below 20%) suggested that both the 0406 subunit and AbpA constituted distinct families within the TasA superfamily (Figure 6b). Additionally, our searches uncovered subdomain-level matches to archaeal flagellin subunits and bacterial type I filament-forming proteins, such as CupE1, FimA, and PapA (Figure 6a), all of which adopted the immunoglobulin-like (Ig) fold. Despite the low sequence similarity, the core structures of these proteins displayed a remarkably conserved topology, hinting at a potential deep homologous relationship between the TasA-related (e.g., CanX, Abp, 0406 thread) and FimA-related proteins (e.g., CupE1, PapA). These findings underscore the evolutionary diversity of cell surface filaments formed through donor strand complementation and suggested a shared structural basis underlying their assembly across different domains of life.

While type IV pili, including the archaeal flagellins, are widespread across archaea, the distribution of TasA-like filaments in archaea remains less well understood. To address this, we used HMMer to search for homologs of bacterial TasA, AbpX, AbpA, CanX, and the 0406-thread subunit in representative archaeal genomes from the Genome Taxonomy Database (GTDB). Our searches revealed that homologs of AbpA, CanX, and the 0406 filament are primarily restricted to the *Thermoproteota* phylum, whereas homologs of AbpX and bacterial TasA are found extensively across several archaeal phyla (Figure 6b). Interestingly, many archaeal species possess two or more of these TasA-like filaments. For example, *V. distributa* DSM 14429 has homologs of CanX (WP_013336327.1), AbpA (WP_148678262.1), and 0406 thread subunit (WP_013336193.1, WP_013336196.1), highlighting the potential diversity and functional versatility of these proteins. These findings demonstrate that TasA-like filaments are far more widespread in archaea than previously recognized, suggesting that many archaeal species may have the capacity to form biofilms. Furthermore, the widespread occurrence of TasA-like proteins in both bacteria and archaea suggested that their last common ancestor already possessed a TasA-like system, which may have been involved in forming cell surface filaments.

## Discussion

Donor strand complementation underlies the polymerization of several classes of bacterial^3–7,9,39^ and archaeal^10,11^ pili that are involved in cellular adhesion and biofilm formation. In contrast to the cannulae, proper assembly of these pili can require specialized cellular machinery such as the chaperone-usher (CU) secretion system.^57,58^ In CU pili,^57,59^ donor strand exchange^60^ leads to the polymerization of Ig-like pilin monomers into extracellular filaments that display an elastomeric or super-elastomeric mechanical response.^3–7,61^ Our results suggested that a similar process of donor strand complementation resulted in polymerization of cannula-like protein monomers into filaments that displayed high mechanical stiffness and thermal stability.^32,35^ Moreover, *in vitro* polymerization of recombinant cannula-like monomers resulted in filaments that preserved the critical interfacial interactions between protomers that were observed in the structure of *ex vivo* cannulae. In comparison, *in vitro* polymerization of CU-type pilins was kinetically slow in absence of the usher catalyst and resulted in pili of limited length and lower thermodynamic stability.^58,62^ In the assembly of CU pili, the chaperone stalls subunits in a higher energy folding intermediate, whose collapse during donor-strand-exchange at the usher drives pilus assembly and concomitant translocation.^63–65^ Here, the need to translocate the OM implies the requirement for this two-step, chaperone-assisted assembly principle that builds an intrinsic free energy gradient across the cell envelope.^65^ In type V pili, which polymerize through an alternative mechanism of donor-strand complementation, subunits reach the surface as lipoproteins and require a proteolytic release of the N-terminal lipid anchor to activate donor strand exchange.^39,66^ In the archaeal cell envelope, however, subunits reach the cell surface directly through the general secretory pathway. Accordingly, unlike most reported protein filaments derived from donor strand complementation, the assembly of the cannulae does not require the presence of a dedicated membrane-bound translocation/assembly platform^67^ or accessory proteins such as chaperones^57^ or proteases^39,66^ to mediate polymerization. Bioinformatic analyses indicated that cannula-like proteins occurred as a divergent family within the larger TasA superfamily of biofilm matrix proteins (Figure 6a, b). TasA-like extracellular filaments, such as the archaeal bundling pili *Pyrobaculum calidifontis* AbpA and *Pyrodictium abyssi* AbpX described here, are limited to monoderm cell envelope architectures of Gram-positive bacteria and archaea.^10^ It is notable that cannulae and ABP polymeric filaments–both derived from donor strand complementation but displaying distinct structures–were co-expressed simultaneously on the cell surface of *P. abyssi* presumably through independent secretory pathways.

The presence of calcium ion binding sites distinguished cannulae from other extracellular filaments that arose from donor strand complementation. Our results suggested that calcium ion coordination primed donor strand complementation and promoted the *in vitro* polymerization of CanA and Hyper2 monomers into cannula-like filaments (Figure 3). In addition, calcium ion coordination stabilized the resultant assemblies through formation of a network of axial and radial interactions between structural subunits in both the *ex vivo* and *in vitro* cannulae. Coordination of calcium has been previously reported to induce structural transitions in proteins secreted at prokaryotic cell surfaces. These transitions have been attributed to the significant Ca^2+^ ion concentration gradient (∼10^4^)^68^ between the extracellular and intracellular environment. Calcium-dependent structural transitions have been studied extensively in extracellular filaments such as the repeats-in-toxin adhesion proteins (RTX adhesins),^69^ which are megadalton proteins that are translocated through the OM of Gram-negative bacteria via the Type I secretion system (T1SS).^70^ RTX adhesins can consist of >100 copies of tandemly repeated bacterial Ig-like (BIg) protein domains capped with a C-terminal RTX β-roll domain. Since the secretion signal is located at the C-terminus of the protein, translocation cannot proceed until completion of protein synthesis, which requires that the nascent polypeptide chains are maintained in an unaggregated state that is permissive for translocation through the T1SS translocon. Biophysical analyses of pathogenic^71^ and non-pathogenic^72,73^ RTX adhesins have demonstrated that calcium ion binding plays several critical roles for this class of proteins including facilitation of transport across the OM and induction of folding in the extracellular environment.^74,75^ RTX adhesin proteins are proposed to be maintained in a fully or partially unfolded state at low intracellular calcium ion concentration prior to secretion through the translocon.^75–77^ In the extracellular environment, the higher calcium ion concentration initiates folding of the BIg protein domains, essentially serving as a “chemical foldase”.^70^ Crystallographic analyses of single or tandemly repeated RTX BIg domains provided evidence that calcium binding not only stabilized the internal structure of individual proteins but also structured flexible loops between domains, causing a rigidification of the structure that enabled the filament to extend outward from the cellular surface.^71,73,75,76,78^ This process is like that which we proposed for polymerization of the cannula-like protein from a comparison of the CanA monomer and protomer structures, in which calcium binding induced structural transitions within loops 34-40 and 163-172 that released the acceptor groove for donor strand complementation. Likewise, the calcium ion gradient has been demonstrated to induce self-assembly of bacterial S-layers.^79–81^ Binding of calcium ion to interfacial loops of these cell surface structures induced a structural transition into their assembly-competent conformation and promoted adoption of the correct quaternary structure. Like the RTX adhesins, calcium-induced structural transitions were proposed to provide a control mechanism to prevent premature assembly of nascent subunits in the cytoplasmic environment, where calcium is held at low nanomolar concentrations.^79,80^

The large lumen (∼18 nm) of the cannula filaments provides an attractive target for the development of functional nanomaterials that can encapsulate and release large substrates but also begs the question of the native function of cannulae. CryoET analysis of *P. abyssi* TAG11 cannulae provided evidence for the presence of density within the lumen, which led to a proposed role in substrate transport.^38^ Our structural analysis of the *ex vivo* cannulae indicated that they could accommodate filamentous cargo within the lumen, which in several instances was observed to be helically wound at the interior surface (Figure 1d). The thin diameter of the filamentous cargo (∼2 nm) and its susceptibility to DNase I treatment suggested the presence of DNA in the lumen, although the structural analysis was inconclusive regarding the cargo identity. Extracellular filaments have been previously demonstrated to mediate horizontal gene transfer (HGT) in prokaryotes through transport of DNA between cells. The transport mechanism proposed in archaea involves the synthesis of a thin conjugative pilus (∼ 7-9 nm outer diameter) through which DNA uptake can occur from a donor cell. We wondered if cannulae could serve as an alternative conduit that could mediate HGT through cross-cellular DNA transport. Conjugative DNA uptake in archaea involves the Ced/Ted DNA import systems,^19,82^ which are encoded on the archaeal chromosome and are homologous to the prokaryotic type IV secretion system (T4SS).^83^ Could cannulae replace the more widespread conjugative DNA uptake process in *Pyrodictiaceae*? To answer this question, we interrogated for the presence of Ced/Ted-related proteins among the family *Pyrodictiaceae*. Pilin protein sequences *Aeropyrum pernix* CedA1 (WP_010865579) and *Pyrobaculum calidifontis* TedF (WP_011849449) were employed to query the genomes of the family *Pyrodictiaceae* in a PSI-BLAST search. The TedF query resulted in no positive hits, which was not surprising given that the families reside in different orders. In contrast, the *A. pernix* CedA1sequence, belonging to the same order *Desulfurococcales*, returned multiple hits at E-values (10^-24^-10^-27^) that were suggestive of high sequence homology. The recovered CedA1 homologues encompassed all members of the family *Pyrodictiaceae* for which genomic information was available as well as MAG data from uncultured *Pyrodictium* and *Hyperthermus* species. While cannulae may play a role in horizontal gene transfer through DNA transport, the conservation of an endogenous Ced DNA import machinery suggested that a cannula-mediated mechanism must be complementary to and potentially independent from HGT through the Ced conjugative uptake pathway.

We speculate that *Pyrodictiaceae* family members that express cannulae may display a clonal form of multicellular organization that is distinct from what is observed in most other microbial communities. These hyperthermophilic archaea, thriving in deep-sea hydrothermal vents at elevated temperature (∼100°C), form intricate colonies with individual cells linked by cannulae, in which the biofilm integrity may be further supported by other matrix proteins like AbpX. The cannulae can potentially enable physical and chemical communication (including possible DNA exchange) between cells, creating a structure somewhat akin to a tissue. Interconnected cells form a mesh-like network that, while not truly multicellular in the sense of differentiated cells in animals or plants, represents a cooperative and coordinated community. This network likely enhances resilience under extreme conditions, allowing for resource sharing (e.g., for DNA repair) and a robustness to environmental stress—characteristics often associated with the early stages of multicellularity.

## Materials and Methods

Luria-Bertani (LB) medium was purchased from IBI Scientific (Dubuque, IA). Isopropyl β-D-1- thiogalactopyranoside (IPTG) was purchased from GoldBio (St. Louis, MO). Chicken egg white lysozyme was purchased from Research Products International (RPI) (Prospect, IL). Benzonase nuclease was purchased from Merck KGaA (Darmstadt, Germany). Protease Inhibitor Cocktail Set V was purchased from Calbiochem (San Diego, CA). Competent cells of *E. coli* strain BL21Gold(DE3) were purchased from Agilent Technologies (Cedar Creek, TX). Precision Plus Protein™ Kaleidoscope™ Pre-stained Protein Standards were purchased from Bio-Rad (Hercules, CA). Custom gene synthesis was performed by ATUM (Newark, CA). Carbon-coated copper grids (200 mesh) and C-flat cryo-EM grids were purchased from Electron Microscopy Sciences (Hatfield, PA). Lacey carbon cryo grids and 2% (w/w) solutions of methylamine vanadate (Nano-Van) and methylamine tungstate (Nano-W) negative stains were purchased from Ted Pella (Redding, CA). All other chemical reagents were purchased from either VWR (Radnor, PA) or Sigma-Aldrich Chemical Co. (St. Louis, MO) and used without further purification.

### Protein expression and purification

Codon-optimized genes corresponding to the expression cassettes of CanA and Hyper2 (Figure S9) were purchased from ATUM (Newark, CA) as sequence-verified clones in plasmid pD451-SR (Kan^R^). The synthetic genes were under control of a T7 promoter with a strong ribosome binding site in high copy number plasmids (pUC *ori*). Purified plasmids encoding the respective proteins were transformed into *E. coli* strain BL21Gold(DE3). Single colonies of the respective transformants were inoculated into sterile Luria-Bertani (LB) medium (4 mL) supplemented with kanamycin (100 μg/mL) and incubated at 37 °C for 12 h. The pre-cultures were used to inoculate sterile LB media (200 mL) in a shake flask supplemented with kanamycin to a final concentration of 100 μg/mL. Bacterial cultures were incubated on a rotary shaker at 37 °C until the OD_600_ reached a value of ca. 0.4 absorption units (AU). Protein expression was induced through addition of isopropyl β-D-1-thiogalactopyranoside (IPTG) to a final concentration of 1 mM and the bacterial cultures were incubated with shaking for an additional 4 h at 30 °C. Cells were isolated from the expression culture through centrifugation at 9100*xg* for 3 min and resuspended in 2 mL of low-salt buffer (50 mM Tris-HCl pH 7.5, 80 mM NaCl, 9% glycerol). The following reagents were added to the cell lysate: hen egg-white lysozyme (final concentration of 1.25 mg/mL), Benzonase nuclease (250 units), MgCl_2_ (final concentration of 1 mM), and 100x protease inhibitor cocktail (to 1x). The cell lysate was incubated with shaking at 30 °C for 1 h. The clarified cell lysate was dialyzed against low-salt buffer containing EDTA (final concentration of 5 mM). After several changes in this high EDTA buffer, the protein was dialyzed in a low-EDTA version of the same buffer (0.01 mM EDTA) with several changes. The dialysate was heated to 80 °C for 20 min to denature the heat-labile proteins of the host bacterium and was subsequently cooled to ambient temperature. The resultant solution was centrifuged at 14000xg for 10 min at ambient temperature. The clarified supernatant fraction was dialyzed overnight against low-salt buffer in a membrane having MWCO of 10,000 Da for CanA and 7,000 Da for Hyper 2. The protein concentration of the dialysate was determined spectrophotometrically using a Nanodrop spectrophotometer. Purified protein samples (4 mL) in low-salt buffer achieved a concentration of ca. 400 µM after purification. Protein purity was assessed using SDS-PAGE gel electrophoresis and electrospray-ionization mass spectrometry.

### Mass Spectrometry

Protein samples were dialyzed extensively against deionized water to remove salt prior to mass spectrometric analysis. Mass spectra were acquired on a Thermo Exactive Plus using a nanoflex source with off-line adapter. Solution of peptide (10 μL) were deposited into a New Objective econo picotip and placed in the nanosource using the off-line adapter. A backing pressure was applied using an off-line syringe to assure flow to the tip. A voltage of 1.0 to 2.5 kV was applied to the picotip. Typical settings employed for the analysis were a capillary temperature of 320 °C and an S-lens RF level between 30-80 with an AGC setting of 1×10^6^. The maximum injection time was set to 50 ms. Spectra were taken at 140,000-resolution at an m/z of 200 using Tune software and analyzed with Thermo Freestyle software.

### Analytical Ultracentrifugation

Sedimentation velocity experiments were performed on a Beckman Colter Optima AUC at the Canadian Center for Hydrodynamics at the University of Lethbridge. CanA samples (9 μM or 31 μM) were measured in low-salt buffer; either alone or supplemented with aqueous CaCl_2_ to a final concentration of 1 - 20 mM. All measurements were conducted in epon-charcoal centerpieces fitted with quartz windows and measured in intensity mode at 45,000 RPM and 20 °C. Data were collected at 220 nm for the low concentration CanA (9 μM) samples and 295 nm for the high concentration CanA (31 μM) samples. All data were analyzed with UltraScan III version 4.0.^84^ The data were fitted with an iterative two-dimensional spectrum analysis^85^ to fit time-and radially invariant noise and meniscus positions. Sample analysis was further refined using Monte Carlo analysis,^86^ and diffusion-corrected integral sedimentation coefficient distributions were generated using the enhanced van Holde-Weischet analysis methods.^87^ UltraScan calculated the buffer density and viscosity to be 1.023810 g/cm^3^ and 1.27097 cP, respectively.

### *In vitro* cannulae assembly

Aliquots (100 μL) of the respective proteins (50-100 μM) in low-salt buffer were incubated in the presence of different concentrations of CaCl_2_ and MgCl_2_ (final concentration from 5 – 20 mM in the respective divalent ions). The temperature of the polymerization mixture for CanA was heated in the temperature range from 50-80 °C for 20 min and cooled to ambient temperature. The polymerization mixture of Hyper2 was incubated for 24 h at ambient temperature after addition of Ca^2+^ ion to the desired concentration. Polymerizations samples were screened using negative-stain transmission electron microscopy (TEM) for the presence of fibrillar assemblies.

### Negative-stain transmission electron microscopy

Aliquots (5 µL) of the polymerization mixtures under different assembly conditions were deposited onto 200-mesh carbon-coated copper grids. After 1 min of incubation, excess solution was removed by wicking the sample. An aliquot (5 µL) of negative stain (1% mixture of 50% NanoVan/50% NanoW) was placed on the grid and incubated for 1 min. Excess stain was wicked away and the grids were air-dried. TEM images were collected with a JEOL JEM-1400 transmission electron microscope at an accelerating voltage of 80 kV.

### Cryo-EM sample preparation, data collection and helical reconstruction of *ex vivo* cannulae

An active culture of *Pyrodictium abyssi* strain AV2 (DSMZ 6158) was purchased from DSMZ-German Collection of Microorganisms and Cell Cultures GmbH. *P. abyssi* was cultured in medium 283 (*Pyrodictium* medium) under anaerobic conditions at 98°C in a pressurized vessel with 2 bar over-pressure of sterile 80% H_2_ and 20% CO_2_ gas mixture. Aliquots (100 µl) of pressurized culture were taken using a 1 mL Injekt-F syringe (Braun) fitted with a 23-gauge sterile needle (BD Microlance) by puncturing the rubber fitting. Harvested aliquots were transferred to 1.5 mL Eppendorf tubes effectively breaking the anaerobe storage conditions. To minimize the effects of exposure to oxygen, care was taken to prepare cryoEM grids shortly after sample harvesting. For that, aliquots (3 µl) of the culture were used as is during grid preparation with no further processing to avoid sample processing artefacts.

A high-resolution cryo-EM dataset was collected using Quantifoil™ R2/1 300 copper mesh holey carbon grids. Grids were glow-discharged at 5 mA plasma current for 30 s in an ELMO (Agar Scientific) glow-discharger. A Gatan CP3 cryo-plunger set at −176 °C and relative humidity of 90% was used to prepare the cryo-samples. A sample (3 μL) of the culture was applied on the holey grid and incubated for 60 s. The sample was back-blotted using Whatman type 2 paper for 3 s and plunge-frozen into precooled liquid ethane at −176 °C. High-resolution movies were recorded at 300 kV on a JEOL Cryoarm300 microscope equipped with an in-column Ω energy filter (operated at slit width of 14 eV) automated with SerialEM 3.0.850. The movies were captured with a K3 direct electron detector run in counting mode at a magnification of 60K with a calibrated pixel size of 0.71 Å/pixel, and a total exposure of 60 e/Å^2^ over 60 frames. A total of 2365 movies were collected and imported into cryoSPARC v4.5.3^88^ for further processing. Movies were motion-corrected using Patch Motion Correction and defocus values were determined using Patch CTF. Exposures were curated and segments were picked using the filament tracer and extracted (750k particles) with a box size of 300 x 300 pixels at a pixel resolution of 1.42 Å/pixel. After several rounds of 2D classification (310k particles retained), Fourier-Bessel indexing was used to calculate initial estimates of the helical rise and twist values, which were used in a subsequent helical refinement job. Next, particles were re-extracted at 0.71 Å/pixel (600 x 600 pixel) and used as input for a second round of helical refinement, followed by local and global CTF refinement, and helical refinement. The resulting high-resolution volume and particle stack were used for reference-based motion correction after particle duplicate removal (310k particles) followed by a final helical refinement (2.30 Å global resolution). An initial atomic model was built using ModelAngelo^41^ without providing an input sequence (build_no_seq), after which the model was manually adjusted to the CanX sequence (WP_338251948.1) and real-space refined in Coot.^89^ A final round of refinement was performed using Phenix^90,91^) and figures were created using ChimeraX.^92^ Map and model statistics are found in Supplementary Table 2.

### Cryo-EM imaging and helical reconstruction of *in vitro* assembled cannulae

The CanA and Hyper2 samples (ca. 3-3.5 μL) were applied to glow-discharged lacey carbon grids and plunge frozen using an EM GP Plunge Freezer (Leica). Grids were positively glow-discharged using amylamine for the Hyper2 sample. The cryo-EM movies were collected on a 300 keV Titan Krios with a K3 camera (University of Virginia) at 1.08 Å/pixel and a total dose of ca. 50 e/Å^2^. First, motion corrections and CTF estimations were done in cryoSPARC^88,93,94^. Next, particles were automatically picked by “Filament Tracer” with a shift of 13 and 24 pixels for CanA and Hyper2, respectively. All auto-picked particles were subsequently 2D classified through multiple rounds of selection, and all particles in bad 2D averages were discarded. After initial processing, 659,059particles remained for CanA. In contrast to the CanA tubes, the Hyper2 assemblies displayed significant polymorphism including the presence of single-walled and multi-walled tubes. The single-walled tubes constituted populations of tubes that displayed different diameters. 2D classification of the single-walled tubes gave enough particles for 3D reconstruction (95,418 and 140,207) for two distinguishable classes representing tubes having diameters of 240 Å (Hyper2 thin tube) and 270 Å (Hyper2 thick tube), respectively. A range of possible helical symmetries was calculated from the respective averaged power spectra derived from aligned raw particles. To speed up the reconstruction, a smaller subset of particles was used to test all possible helical symmetries by trial and error until amino acid side chains, and the hand of a short α-helix was seen.^49,95^ The resolution of the correct symmetry was estimated by Map:Map FSC, Model:Map FSC, and d_99_.^96^ The final volumes were then sharpened with a negative B-factor,-85 Å^2^,-88 Å^2^ and-89 Å^2^ for CanA, Hyper2 thick tube and Hyper2 thin tube, respectively, which was automatically estimated in cryoSPARC. Maps were further sharpened by EMReady.^97^ FSC calculations were employed to estimate the resolution of the reconstructions and are reported in Supplementary Figure 14, 17, and Supplementary Table S2.

### Model Building

The absolute helical hand of the maps of CanA and Hyper2 tubes was determined from the hand of an α-helix. The hand assignment also agreed with the AlphaFold2^44^ prediction of CanA and Hyper2. The cryo-EM map corresponding to a single subunit of CanA and Hyper2 was segmented in Chimera.^98^ The AlphaFold predictions of CanA and Hyper2 were used as the starting model and docked into the cryoEM maps. The docked models were refined by iterative cycles of manual refinement using Coot^89^ and automatic refinement using PHENIX^99^ and ISOLDE^100^ as needed to reach the reasonable map-to-model agreement and model geometry. Density for eight (1-8) N-terminal residues and four (149-152) C-terminal residues of Hyper2 was not resolved in the map and thus these residues were left unmodeled. MolProbity^101^ was used to evaluate the quality of the filament model. The refinement statistics are shown in Supplementary Table 2.

### Crystallization and data collection

Crystallization of the mature CanA protein was concentrated to 10 mg/ml then put into crystallization trials. All crystals were grown by sitting drop vapor diffusion at 20°C using a protein to reservoir volume ratio of 1:1 with total drop volumes of 0.2 μL. Crystals of CanA were grown using a crystallization solution containing 100 mM MgCl2, 30% PEG 4K, 3% Xylitol, and 100 mM Tris pH 8.5. All crystals were flash frozen in liquid nitrogen after a short soak in the appropriate crystallization buffers supplemented with 20 - 25% ethylene glycol. Data for CanA were collected at the beamline 19-ID NYX at the National Synchrotron Light source II (NSLS II), Brookhaven National Labs. All data was indexed, merged, and scaled using HKL2000 then converted to structure factors using CCP4.

### Crystal structure determination and refinement

The CanA crystal structure was solved by molecular replacement using the program Phaser. Molecular replacement calculations were performed using the coordinates of the CanA monomeric subunit from the CanA cryoEM structure (PDB: 7UII) as the search model. The resulting phase information from molecular replacement was used for manual model fitting of the CanA structure using the graphics program COOT.^89^ <u>S</u>tructural refinement of the CanA coordinates were performed using the PHENIX package.^90^ During refinement a cross-validation test set was created from a random 5% of the reflections. Data collection and refinement statistics are listed in Supplementary Table 3. Molecular graphics of the CanA crystal structure were prepared using the PyMOL Molecular Graphics System (Schrodinger) (DeLano Scientific LLC, Palo Alto, CA) or ChimeraX.^92^

### Bioinformatic analysis

All sequence similarity searches were performed using the HMMER webserver^46^ against the UniProtKB database and a local installation of HMMER v3.4^102^ against the non-redundant (nr) protein sequence database. Additionally, HHsearch within the MPI Bioinformatics Toolkit^103^ was utilized to detect distant homologs.

For pairwise sequence comparisons of representative archaeal and bacterial cell-surface filament-forming proteins (Figure 6a), we constructed multiple sequence alignments by running three iterations of HHblits^104^ against the UniClust30 database.^105^ Secondary structure information was incorporated into these alignments using the addss.pl script from HH-suite3. Profile hidden Markov models (HMMs) were then generated using hhmake, and context-specific transitions were calculated using cstranslate. A custom-built profile HMM database was built as outlined in the HH- suite3 user guide. To obtain pairwise HHsearch probabilities, individual alignments were searched against this custom-built profile HMM database using default parameters. Figure 6a includes the following proteins: CanX from *Pyrodictium abyssi* (WP_338251948.1) and *Hyperthermus* sp. (RUM47266.1); CanX-like from *Saccharolobus solfataricus* (WP_009990924.1, WP_009991716.1, WP_029552507.1); AbpX from *P. abyssi* (WP_338249486.1); AbpA from *Pyrobaculum calidifontis* (A3MUL8); 0406 thread subunit from *Sulfolobus acidocaldarius* (Q4JBK8); TasA from *Archaeoglobus fulgidus* (A0A075WE38), *Methanosarcina barkeri* (A0A0E3QZX2), and *Bacillus subtilis* (P54507); CalY1 from *B. subtilis* (AKB91166); Bacterial type I pilins from *Escherichia coli* (PapA - P04127, FimA1-P04128), *Acinetobacter baumannii* (CsuA/B - A0A6F8TDQ5), and *Pseudomonas aeruginosa* (CupE1-P04128); Archaeal flagellin subunit from *Methanospirillum hungatei* (Q2FUM4), *Methanococcus maripaludis* (Q6LWP3), and *Pyrococcus furiosus* (A0A0B4ZYM1); Type IV pilins from *S. solfataricus* (A0A157T322, UpsA - Q7LXX9), *Sulfolobus islandicus* (M9UD72), and *S. acidocaldarius* (Q4J6I2); and Bacterial type IV pilins from *Thermus thermophilus* (Q72GL2), *P. aeruginosa* (PSEAIP02973), and *Neisseria gonorrhoeae* (P02974).

To investigate the prevalence of TasA-like filaments in archaea (Figure 6b), we searched for homologs of TasA, AbpX, AbpA, the 0406-thread subunit, and CanX-like proteins in archaeal proteomes within the Genome Taxonomy Database (GTDB; release R220)^106^ using HMMER. First, we built multiple sequence alignments by running three iterations of jackhmmer against the nr70_arc database, a filtered version of the nr protein sequence database containing only archaeal sequences and with a maximum pairwise sequence identity of 70%. For TasA, we used the pre-existing alignment (PF12389) from the Pfam database.^107^ The resulting alignments were then converted into profile HMMs using hmmbuild. To detect homologs of each query protein in the GTDB archaeal proteomes, we performed searches with hmmsearch using a stringent E-value threshold (-E) of 1e-10. We visualized the presence of these homologs in a GTDB archaeal phylogenetic tree using iTOL.^108^ For this, we converted the GTDB archaeal trees into iTOL-compatible format using the convert_to_itol method from GTDB-tk v2.4.0+.^109^ For signal peptide prediction, we used DeepTMHMM,^110^ and for obtaining structural models, we used a local installation of AlphaFold2,^44^ the AlphaFold3 webserver,^48^ as well as the AlphaFold Protein Structure Database.^111^

## Acknowledgments

High-resolution cryo-EM imaging was conducted at the Molecular Electron Microscopy Core facility at the University of Virginia, which is supported by the School of Medicine and built with NIH grant G20-RR31199. In addition, the Titan Krios (SIG S10-RR025067) and K3/GIF (U24-GM116790) were purchased in part or in full using the designated NIH grants. This study was supported by the Robert P. Apkarian Integrated Electron Microscopy Core (IEMC) at Emory University, which is subsidized by the School of Medicine and Emory College of Arts and Sciences. Additional support was provided by the Georgia Clinical & Translational Science Alliance of the National Institutes of Health under award number UL1TR000454. Negative stain TEM images gathered on a Hitachi HT7700 120 kV TEM at Emory University, which was supported by the Georgia Clinical and Translational Science Alliance under award number UL1TR002378. Use of the NYX beamline 19-ID at the National Synchrotron Light Source II was supported by the New York Structural Biology Center. This research used resources of the National Synchrotron Light Source II, a U.S. Department of Energy (DOE) Office of Science User Facility operated for the DOE Office of Science by Brookhaven National Laboratory under Contract No. DE-SC0012704. NYX detector instrumentation was supported by grant S10OD030394 through the Office of the Director, National Institutes of Health. The AUC studies were supported by the Canada 150 Research Chairs program C150-2017-00015, the National Institutes of Health grant 1R01GM120600, and the Canadian Natural Science and Engineering Research Council Discovery Grant DG-RGPIN-2019-05637 to B.D. The Canadian Center for Hydrodynamics is funded by the Canada Foundation for Innovation grant CFI-37589 (BD). UltraScan supercomputer calculations were supported through NSF/XSEDE grant TG-MCB070039N to B.D. This research was supported by grants from NSF (2003962) to V.P.C, NIH (GM138756) to F.W, and NIH (GM122510) to E.H.E. The content is solely the responsibility of the authors and does not necessarily reflect the official views of the National Institutes of Health.

## Data Availability

The reconstruction maps were deposited in the Electron Microscopy Data Bank with accession numbers of EMD-51935 (CanX), EMD-26546 (CanA) and EMD-44403 (Hyper2thick tube), and EMD- 44404 (Hyper2-thin tube). The corresponding filament models were deposited in the Protein Data Bank with accession numbers of 9H8B (CanX), 7UII (CanA) and 9BAB (Hyper2-thick tube) and 9BAC (Hyper2-thin tube). The coordinate file and diffraction data for the CanA crystal structure were deposited into the Protein Data Bank with accession code 9DLO.

## References

1 Egelman, E. H. Cryo-EM of bacterial pili and archaeal flagellar filaments. Current Opinion in Structural Biology 46, 31–37 (2017). 10.1016/j.sbi.2017.05.012

2 Kreutzberger, M. A. B. et al. Convergent evolution in the supercoiling of prokaryotic flagellar filaments. Cell 185, 3487–3500 e3414 (2022). 10.1016/j.cell.2022.08.009

3 Hospenthal, M. K. et al. Structure of a Chaperone-Usher Pilus Reveals the Molecular Basis of Rod Uncoiling. Cell 164, 269–278 (2016). 10.1016/j.cell.2015.11.049

4 Spaulding, C. N. et al. Functional role of the type 1 pilus rod structure in mediating host-pathogen interactions. Elife 7 (2018). 10.7554/eLife.31662

5 Pakharukova, N. et al. Archaic chaperone-usher pili self-secrete into superelastic zigzag springs. Nature 609, 335–340 (2022). 10.1038/s41586-022-05095-0

6 Doran, M. H., Baker, J. L., Dahlberg, T., Andersson, M. & Bullitt, E. Three structural solutions for bacterial adhesion pilus stability and superelasticity. Structure 31, 529–540 e527 (2023). 10.1016/j.str.2023.03.005

7 Bohning, J. et al. Architecture of the biofilm-associated archaic Chaperone-Usher pilus CupE from Pseudomonas aeruginosa. PLoS Pathog 19, e1011177 (2023). 10.1371/journal.ppat.1011177

8 Hospenthal, M. K. et al. The Cryoelectron Microscopy Structure of the Type 1 Chaperone-Usher Pilus Rod. Structure 25, 1829–1838 e1824 (2017). 10.1016/j.str.2017.10.004

9 Bohning, J. et al. Donor-strand exchange drives assembly of the TasA scaffold in Bacillus subtilis biofilms. Nat Commun 13, 7082 (2022). 10.1038/s41467-022-34700-z

10 Wang, F., Cvirkaite-Krupovic, V., Krupovic, M. & Egelman, E. H. Archaeal bundling pili of Pyrobaculum calidifontis reveal similarities between archaeal and bacterial biofilms. Proc Natl Acad Sci U S A 119, e2207037119 (2022). 10.1073/pnas.2207037119

11 Gaines, M. C. et al. Electron cryo-microscopy reveals the structure of the archaeal thread filament. Nat Commun 13, 7411 (2022). 10.1038/s41467-022-34652-4

12 Sleutel, M., Pradhan, B., Volkov, A. N. & Remaut, H. Structural analysis and architectural principles of the bacterial amyloid curli. Nat Commun 14, 2822 (2023). 10.1038/s41467-023-38204-2

13 Wang, F. et al. Structure of Microbial Nanowires Reveals Stacked Hemes that Transport Electrons over Micrometers. Cell 177, 361–369.e310 (2019). 10.1016/j.cell.2019.03.029

14 Wang, F. et al. Cryo-EM structure of an extracellular Geobacter OmcE cytochrome filament reveals tetrahaem packing. Nat Microbiol 7, 1291–1300 (2022). 10.1038/s41564-022-01159-z

15 Wang, F. et al. Structure of Geobacter OmcZ filaments suggests extracellular cytochrome polymers evolved independently multiple times. Elife 11 (2022). 10.7554/eLife.81551

16 Baquero, D. P. et al. Extracellular cytochrome nanowires appear to be ubiquitous in prokaryotes. Cell 186, 2853–2864 e2858 (2023). 10.1016/j.cell.2023.05.012

17 Costa, T. R. D. et al. Structure of the Bacterial Sex F Pilus Reveals an Assembly of a Stoichiometric Protein-Phospholipid Complex. Cell 166, 1436–1444.e1410 (2016). 10.1016/j.cell.2016.08.025

18 Zheng, W. et al. Cryoelectron-Microscopic Structure of the pKpQIL Conjugative Pili from Carbapenem-Resistant Klebsiella pneumoniae. Structure 28, 1321–1328 e1322 (2020). 10.1016/j.str.2020.08.010

19 Beltran, L. C. et al. Archaeal DNA-import apparatus is homologous to bacterial conjugation machinery. Nat Commun 14, 666 (2023). 10.1038/s41467-023-36349-8

20 Loquet, A. et al. Atomic model of the type III secretion system needle. Nature 486, 276–279 (2012). 10.1038/nature11079

21 Flacht, L. et al. Integrative structural analysis of the type III secretion system needle complex from Shigella flexneri. Protein Sci 32, e4595 (2023). 10.1002/pro.4595

22 Hvorecny, K. L. & Kollman, J. M. Greater than the sum of parts: Mechanisms of metabolic regulation by enzyme filaments. Curr Opin Struct Biol 79, 102530 (2023). 10.1016/j.sbi.2023.102530

23 Zhu, J. et al. Protein Assembly by Design. Chem Rev 121, 13701–13796 (2021). 10.1021/acs.chemrev.1c00308

24 Egelman, E. H. Three-dimensional reconstruction of helical polymers. Archives of Biochemistry and Biophysics 581, 54–58 (2015). 10.1016/j.abb.2015.04.004

25 Wang, F., Gnewou, O., Solemanifar, A., Conticello, V. P. & Egelman, E. H. Cryo-EM of Helical Polymers. Chem Rev (2022). 10.1021/acs.chemrev.1c00753

26 Miller, J. G., Hughes, S. A., Modlin, C. & Conticello, V. P. Structures of synthetic helical filaments and tubes based on peptide and peptido-mimetic polymers. Q Rev Biophys, 1–103 (2022). 10.1017/S0033583522000014

27 Fulton, D. A., Dura, G. & Peters, D. T. The polymer and materials science of the bacterial fimbriae Caf1. Biomater Sci 11, 7229–7246 (2023). 10.1039/d3bm01075a

28 Nguyen, P. Q., Courchesne, N. D., Duraj-Thatte, A., Praveschotinunt, P. & Joshi, N. S. Engineered Living Materials: Prospects and Challenges for Using Biological Systems to Direct the Assembly of Smart Materials. Adv Mater 30, e1704847 (2018). 10.1002/adma.201704847

29 Stetter, K. O. Ultrathin mycelia-forming organisms from submarine volcanic areas having an optimum growth temperature of 105 °C. Nature 300, 258–260 (1982). 10.1038/300258a0

30 Stetter, K. O., Konig, H. & Stackebrandt, E. Pyrodictium gen. nov., a New Genus of Submarine Disc-Shaped Sulphur Reducing Archaebacteria Growing Optimally at 105 degrees C. Syst Appl Microbiol 4, 535–551 (1983). 10.1016/S0723-2020(83)80011-3

31 König, H., Messner, P. & Stetter, K. O. The fine structure of the fibers of Pyrodictium occultum. FEMS Microbiology Letters 49, 207–212 (1988). 10.1111/j.1574-6968.1988.tb02717.x

32 Rieger, G., Rachel, R., Hermann, R. & Stetter, K. O. Ultrastructure of the Hyperthermophilic Archaeon Pyrodictium abyssi. J Struct Biol 115, 78–87 (1995). 10.1006/jsbi.1995.1032

33 Pley, U. et al. Pyrodictium abyssi sp. nov. Represents a Novel Heterotrophic Marine Archaeal Hyperthermophile Growing at 110°C. Systematic and Applied Microbiology 14, 245–253 (1991). 10.1016/S0723-2020(11)80376-0

34 Lin, T. J. et al. Pyrodictium delaneyi sp. nov., a hyperthermophilic autotrophic archaeon that reduces Fe(III) oxide and nitrate. Int J Syst Evol Microbiol 66, 3372–3376 (2016). 10.1099/ijsem.0.001201

35 Barton, N. R. O. d., E.; Short, R.; Frey, G.; Weiner, D.; Robertson, D. E.; Briggs, S.; Zorner, P. Chimeric Cannulae Proteins, Nucleic Acids Encoding Them And Methods For Making And Using Them. United States patent (2008).

36 Rieger, G., Müller, K., Hermann, R., Stetter, K. O. & Rachel, R. Cultivation of hyperthermophilic archaea in capillary tubes resulting in improved preservation of fine structures. Archives of Microbiology 168, 373–379 (1997). 10.1007/s002030050511

37 Horn, C., Paulmann, B., Kerlen, G., Junker, N. & Huber, H. In vivo observation of cell division of anaerobic hyperthermophiles by using a high-intensity dark-field microscope. J Bacteriol 181, 5114–5118 (1999). 10.1128/JB.181.16.5114-5118.1999

38 Nickell, S., Hegerl, R., Baumeister, W. & Rachel, R. Pyrodictium cannulae enter the periplasmic space but do not enter the cytoplasm, as revealed by cryo-electron tomography. J Struct Biol 141, 34–42 (2003). 10.1016/s1047-8477(02)00581-6

39 Shibata, S. et al. Structure of polymerized type V pilin reveals assembly mechanism involving protease-mediated strand exchange. Nat Microbiol 5, 830–837 (2020). 10.1038/s41564-020-0705-1

40 Wang, T. et al. CryoSeek: A strategy for bioentity discovery using cryoelectron microscopy. Proc Natl Acad Sci U S A 121, e2417046121 (2024). https://doi.org:doi:10.1073/pnas.2417046121

41 Jamali, K. et al. Automated model building and protein identification in cryo-EM maps. Nature 628, 450–457 (2024). 10.1038/s41586-024-07215-4

42 Kreitner, R. et al. Complete sequential assignment and secondary structure prediction of the cannulae forming protein CanA from the hyperthermophilic archaeon Pyrodictium abyssi. Biomol NMR Assign 14, 141–146 (2020). 10.1007/s12104-020-09934-x

43 Jarrell, K. F. et al. N-linked glycosylation in Archaea: a structural, functional, and genetic analysis. Microbiol Mol Biol Rev 78, 304–341 (2014). 10.1128/MMBR.00052-13

44 Jumper, J. et al. Highly accurate protein structure prediction with AlphaFold. Nature 596, 583–589 (2021). 10.1038/s41586-021-03819-2

45 Beltran, L. et al. The mating pilus of E. coli pED208 acts as a conduit for ssDNA during horizontal gene transfer. mBio 15, e0285723 (2024). 10.1128/mbio.02857-23

46 Potter, S. C. et al. HMMER web server: 2018 update. Nucleic Acids Res 46, W200–W204 (2018). 10.1093/nar/gky448

47 van Kempen, M. et al. Fast and accurate protein structure search with Foldseek. Nat Biotechnol 42, 243–246 (2024). 10.1038/s41587-023-01773-0

48 Abramson, J. et al. Accurate structure prediction of biomolecular interactions with AlphaFold 3. Nature 630, 493–500 (2024). 10.1038/s41586-024-07487-w

49 Egelman, E. H. Reconstruction of helical filaments and tubes. Methods Enzymol 482, 167–183 (2010). 10.1016/s0076-6879(10)82006-3

50 Egelman, E. H. The iterative helical real space reconstruction method: surmounting the problems posed by real polymers. J Struct Biol 157, 83–94 (2007). 10.1016/j.jsb.2006.05.015

51 Krissinel, E. & Henrick, K. Inference of macromolecular assemblies from crystalline state. J Mol Biol 372, 774–797 (2007). 10.1016/j.jmb.2007.05.022

52 Wang, C. L., Leavis, P. C. & Gergely, J. Kinetic studies show that Ca2+ and Tb3+ have different binding preferences toward the four Ca2+-binding sites of calmodulin. Biochemistry 23, 6410–6415 (1984). 10.1021/bi00321a020

53 UniProt, C. UniProt: the Universal Protein Knowledgebase in 2023. Nucleic Acids Res 51, D523–D531 (2023). 10.1093/nar/gkac1052

54 Puorger, C., Vetsch, M., Wider, G. & Glockshuber, R. Structure, folding and stability of FimA, the main structural subunit of type 1 pili from uropathogenic Escherichia coli strains. J Mol Biol 412, 520–535 (2011). 10.1016/j.jmb.2011.07.044

55 Zyla, D., Echeverria, B. & Glockshuber, R. Donor strand sequence, rather than donor strand orientation, determines the stability and non-equilibrium folding of the type 1 pilus subunit FimA. J Biol Chem 295, 12437–12448 (2020). 10.1074/jbc.RA120.014324

56 Evans, R. et al. Protein complex prediction with AlphaFold-Multimer. bioRxiv, 2021.2010.2004.463034 (2022). 10.1101/2021.10.04.463034

57 Waksman, G. Structural and Molecular Biology of a Protein-Polymerizing Nanomachine for Pilus Biogenesis. J Mol Biol 429, 2654–2666 (2017). 10.1016/j.jmb.2017.05.016

58 Zyla, D. S. et al. The assembly platform FimD is required to obtain the most stable quaternary structure of type 1 pili. Nat Commun 15, 3032 (2024). 10.1038/s41467-024-47212-9

59 Sauer, F. G. et al. Structural basis of chaperone function and pilus biogenesis. Science 285, 1058–1061 (1999). 10.1126/science.285.5430.1058

60 Remaut, H. et al. Donor-strand exchange in chaperone-assisted pilus assembly proceeds through a concerted beta strand displacement mechanism. Mol Cell 22, 831–842 (2006). 10.1016/j.molcel.2006.05.033

61 Alonso-Caballero, A. et al. Mechanical architecture and folding of E. coli type 1 pilus domains. Nat Commun 9, 2758 (2018). 10.1038/s41467-018-05107-6

62 Nishiyama, M., Ishikawa, T., Rechsteiner, H. & Glockshuber, R. Reconstitution of pilus assembly reveals a bacterial outer membrane catalyst. Science 320, 376–379 (2008). 10.1126/science.1154994

63 Sauer, F. G., Pinkner, J. S., Waksman, G. & Hultgren, S. J. Chaperone priming of pilus subunits facilitates a topological transition that drives fiber formation. Cell 111, 543–551 (2002). 10.1016/s0092-8674(02)01050-4

64 Zavialov, A. V. et al. Resolving the energy paradox of chaperone/usher-mediated fibre assembly. Biochem J 389, 685–694 (2005). 10.1042/BJ20050426

65 Phan, G. et al. Crystal structure of the FimD usher bound to its cognate FimC-FimH substrate. Nature 474, 49–53 (2011). 10.1038/nature10109

66 Xu, Q. et al. A Distinct Type of Pilus from the Human Microbiome. Cell 165, 690–703 (2016). 10.1016/j.cell.2016.03.016

67 Costa, T. R. D. et al. Secretion systems in Gram-negative bacteria: structural and mechanistic insights. Nature Reviews Microbiology 13, 343–359 (2015). 10.1038/nrmicro3456

68 Bronner, F. Extracellular and intracellular regulation of calcium homeostasis. ScientificWorldJournal 1, 919–925 (2001). 10.1100/tsw.2001.489

69 Guo, S., Vance, T. D. R., Stevens, C. A., Voets, I. & Davies, P. L. RTX Adhesins are Key Bacterial Surface Megaproteins in the Formation of Biofilms. Trends Microbiol 27, 453–467 (2019). 10.1016/j.tim.2018.12.003

70 Spitz, O. et al. Type I Secretion Systems-One Mechanism for All? Microbiol Spectr 7 (2019). 10.1128/microbiolspec.PSIB-0003-2018

71 Griessl, M. H. et al. Structural insight into the giant Ca(2)(+)-binding adhesin SiiE: implications for the adhesion of Salmonella enterica to polarized epithelial cells. Structure 21, 741–752 (2013). 10.1016/j.str.2013.02.020

72 Guo, S. et al. Role of Ca(2)(+) in folding the tandem beta-sandwich extender domains of a bacterial ice-binding adhesin. FEBS J 280, 5919–5932 (2013). 10.1111/febs.12518

73 Vance, T. D. et al. Ca2+-stabilized adhesin helps an Antarctic bacterium reach out and bind ice. Biosci Rep 34 (2014). 10.1042/BSR20140083

74 Bumba, L. et al. Calcium-Driven Folding of RTX Domain beta-Rolls Ratchets Translocation of RTX Proteins through Type I Secretion Ducts. Mol Cell 62, 47–62 (2016). 10.1016/j.molcel.2016.03.018

75 Peters, B. et al. Structural and functional dissection reveals distinct roles of Ca2+-binding sites in the giant adhesin SiiE of Salmonella enterica. PLoS Pathog 13, e1006418 (2017). 10.1371/journal.ppat.1006418

76 Vance, T. D. R., Ye, Q., Conroy, B. & Davies, P. L. Essential role of calcium in extending RTX adhesins to their target. J Struct Biol X 4, 100036 (2020). 10.1016/j.yjsbx.2020.100036

77 Vance, T. D. R., Guo, S., Assaie-Ardakany, S., Conroy, B. & Davies, P. L. Structure and functional analysis of a bacterial adhesin sugar-binding domain. PLoS One 14, e0220045 (2019). 10.1371/journal.pone.0220045

78 Oude Vrielink, A. S., Vance, T. D., de Jong, A. M., Davies, P. L. & Voets, I. K. Unusually high mechanical stability of bacterial adhesin extender domains having calcium clamps. PLoS One 12, e0174682 (2017). 10.1371/journal.pone.0174682

79 Baranova, E. et al. SbsB structure and lattice reconstruction unveil Ca2+ triggered S-layer assembly. Nature 487, 119–122 (2012). 10.1038/nature11155

80 Herdman, M. et al. High-resolution mapping of metal ions reveals principles of surface layer assembly in Caulobacter crescentus cells. Structure 30, 215–228 e215 (2022). 10.1016/j.str.2021.10.012

81 Sogues, A. et al. Structure and function of the EA1 surface layer of Bacillus anthracis. Nat Commun 14, 7051 (2023). 10.1038/s41467-023-42826-x

82 van Wolferen, M., Wagner, A., van der Does, C. & Albers, S. V. The archaeal Ced system imports DNA. Proc Natl Acad Sci U S A 113, 2496–2501 (2016). 10.1073/pnas.1513740113

83 Costa, T. R. D. et al. Type IV secretion systems: Advances in structure, function, and activation. Mol Microbiol 115, 436–452 (2021). 10.1111/mmi.14670

84 Demeler, B. & Gorbet, G. E. in Analytical Ultracentrifugation 119–143 (Springer, 2016).

85 Brookes, E., Cao, W. & Demeler, B. A two-dimensional spectrum analysis for sedimentation velocity experiments of mixtures with heterogeneity in molecular weight and shape. European Biophysics Journal 39, 405–414 (2010).

86 Demeler, B. & Brookes, E. Monte Carlo analysis of sedimentation experiments. Colloid and Polymer Science 286, 129–137 (2008).

87 Demeler, B. & Van Holde, K. E. Sedimentation velocity analysis of highly heterogeneous systems. Analytical biochemistry 335, 279–288 (2004).

88 Punjani, A., Rubinstein, J. L., Fleet, D. J. & Brubaker, M. A. cryoSPARC: algorithms for rapid unsupervised cryo-EM structure determination. Nat Methods 14, 290–296 (2017). 10.1038/nmeth.4169

89 Emsley, P. & Cowtan, K. Coot: model-building tools for molecular graphics. Acta Crystallogr D Biol Crystallogr 60, 2126–2132 (2004). 10.1107/S0907444904019158

90 Adams, P. D. et al. PHENIX: a comprehensive Python-based system for macromolecular structure solution. Acta Crystallogr D Biol Crystallogr 66, 213–221 (2010). doi:10.1107/S0907444909052925

91 Liebschner, D. et al. Macromolecular structure determination using X-rays, neutrons and electrons: recent developments in Phenix. Acta Crystallogr D Struct Biol 75, 861–877 (2019). 10.1107/S2059798319011471

92 Goddard, T. D. et al. UCSF ChimeraX: Meeting modern challenges in visualization and analysis. Protein Sci 27, 14–25 (2018). 10.1002/pro.3235

93 Rohou, A. & Grigorieff, N. CTFFIND4: Fast and accurate defocus estimation from electron micrographs. J Struct Biol 192, 216–221 (2015). 10.1016/j.jsb.2015.08.008

94 Zheng, S. Q. et al. MotionCor2: anisotropic correction of beam-induced motion for improved cryo-electron microscopy. Nat Methods 14, 331–332 (2017). 10.1038/nmeth.4193

95 Egelman, E. H. A robust algorithm for the reconstruction of helical filaments using single-particle methods. Ultramicroscopy 85, 225–234 (2000). 10.1016/s0304-3991(00)00062-0

96 Afonine, P. V. et al. New tools for the analysis and validation of cryo-EM maps and atomic models. Acta Crystallogr D Struct Biol 74, 814–840 (2018). 10.1107/S2059798318009324

97 He, J., Li, T. & Huang, S. Y. Improvement of cryo-EM maps by simultaneous local and non-local deep learning. Nat Commun 14, 3217 (2023). 10.1038/s41467-023-39031-1

98 Pettersen, E. F. et al. UCSF Chimera--a visualization system for exploratory research and analysis. J Comput Chem 25, 1605–1612 (2004). 10.1002/jcc.20084

99 Afonine, P. V. et al. Real-space refinement in PHENIX for cryo-EM and crystallography. Acta Crystallogr D Struct Biol 74, 531–544 (2018). 10.1107/S2059798318006551

100 Croll, T. I. ISOLDE: a physically realistic environment for model building into low-resolution electron-density maps. Acta Crystallogr D Struct Biol 74, 519–530 (2018). 10.1107/S2059798318002425

101 Williams, C. J. et al. MolProbity: More and better reference data for improved all-atom structure validation. Protein Sci 27, 293–315 (2018). 10.1002/pro.3330

102 Eddy, S. R. Accelerated Profile HMM Searches. PLoS Comput Biol 7, e1002195 (2011). 10.1371/journal.pcbi.1002195

103 Zimmermann, L. et al. A Completely Reimplemented MPI Bioinformatics Toolkit with a New HHpred Server at its Core. J Mol Biol 430, 2237–2243 (2018). 10.1016/j.jmb.2017.12.007

104 Steinegger, M. et al. HH-suite3 for fast remote homology detection and deep protein annotation. BMC Bioinformatics 20, 473 (2019). 10.1186/s12859-019-3019-7

105 Mirdita, M. et al. Uniclust databases of clustered and deeply annotated protein sequences and alignments. Nucleic Acids Res 45, D170–D176 (2017). 10.1093/nar/gkw1081

106 Parks, D. H. et al. GTDB: an ongoing census of bacterial and archaeal diversity through a phylogenetically consistent, rank normalized and complete genome-based taxonomy. Nucleic Acids Res 50, D785–D794 (2022). 10.1093/nar/gkab776

107 El-Gebali, S. et al. The Pfam protein families database in 2019. Nucleic Acids Res 47, D427–D432 (2019). 10.1093/nar/gky995

108 Letunic, I. & Bork, P. Interactive Tree of Life (iTOL) v6: recent updates to the phylogenetic tree display and annotation tool. Nucleic Acids Res 52, W78–W82 (2024). 10.1093/nar/gkae268

109 Chaumeil, P. A., Mussig, A. J., Hugenholtz, P. & Parks, D. H. GTDB-Tk v2: memory friendly classification with the genome taxonomy database. Bioinformatics 38, 5315–5316 (2022). 10.1093/bioinformatics/btac672

110 Hallgren, J. et al. DeepTMHMM predicts alpha and beta transmembrane proteins using deep neural networks. bioRxiv, 2022.2004.2008.487609 (2022). 10.1101/2022.04.08.487609

111 Varadi, M. et al. AlphaFold Protein Structure Database in 2024: providing structure coverage for over 214 million protein sequences. Nucleic Acids Res 52, D368–D375 (2024). 10.1093/nar/gkad1011

