## Supplementary Information for "Donor Strand Complementation and Calcium Ion Coordination Drive the Chaperone-free Polymerization of Archaeal Cannulae"

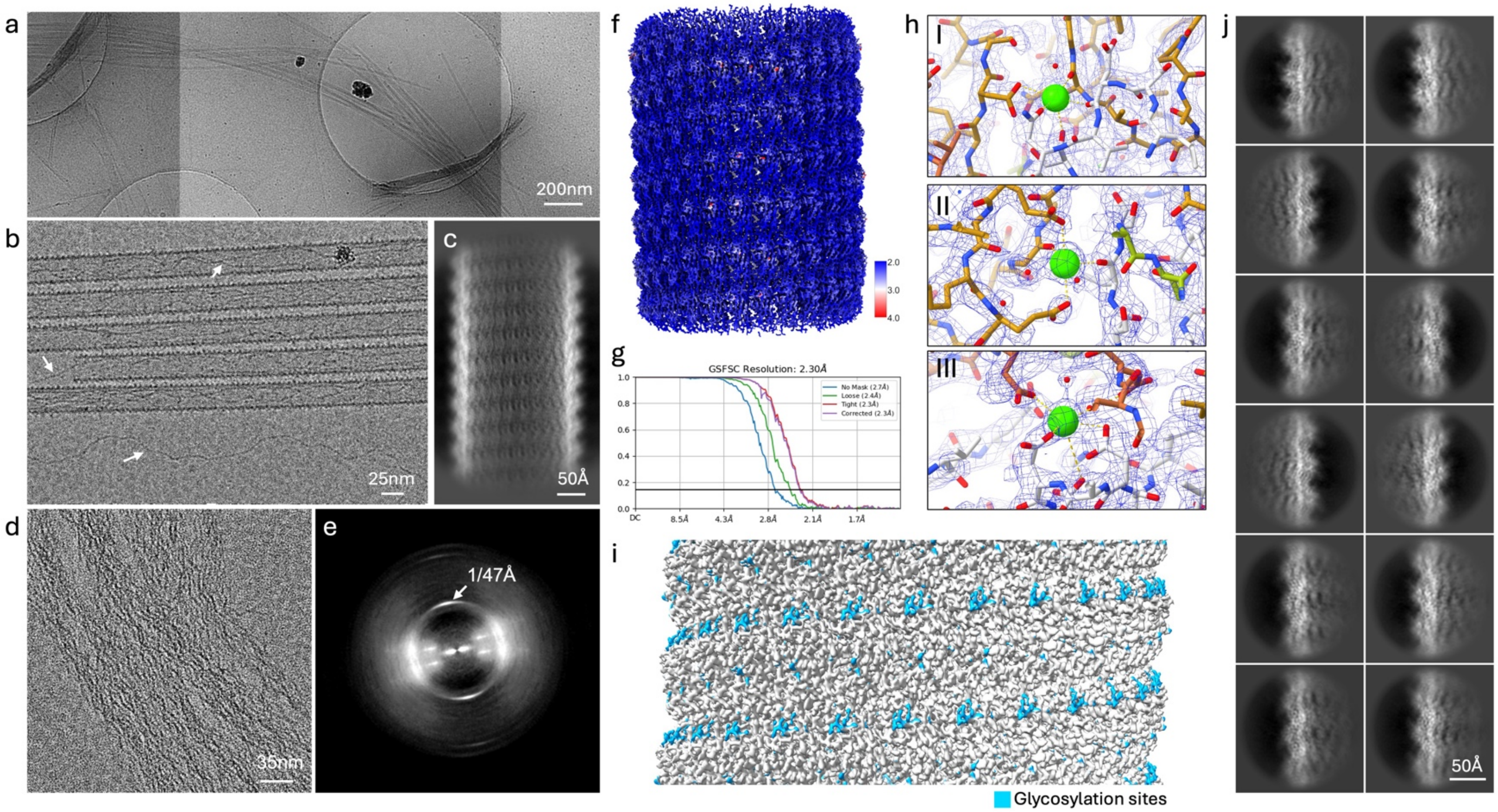


**Supplementary Figure 1: CryoEM observations of *ex vivo* cannulae from *Pyrodictium abyssi*.** Low magnification (2000x) stitched cryoEM micrograph of a plunge frozen active culture of *Pyrodictium abyssi* AV2. Cannulae are present as long (> 5µm) fibers that cluster into larger bundles; (b) High magnification (60.000x) denoised cryoEM micrograph of cannulae fibers that are decorated with 2 nm cargo fibrils; (c) 2D class average: 800x800 pix; 0.7 Å/pix; (d) High magnification (60.000x) denoised cryoEM micrograph of a second type of protein fibers (AbpX) present as super bundles with the corresponding power spectrum in (e) indicating a pitch of 47 Å; (f) *ex vivo* cannulae volume surface colored according local resolution; (f) FSC curve; (h) zoom-ins of the three calcium binding pocket regions with the electrostatic potential map overlaid in blue; (i) glycosylation sites on the cannulae surface; (j) 2D class averages of particles boxed (256x256 pix; 1.4 Å/pix) around the filament edge.

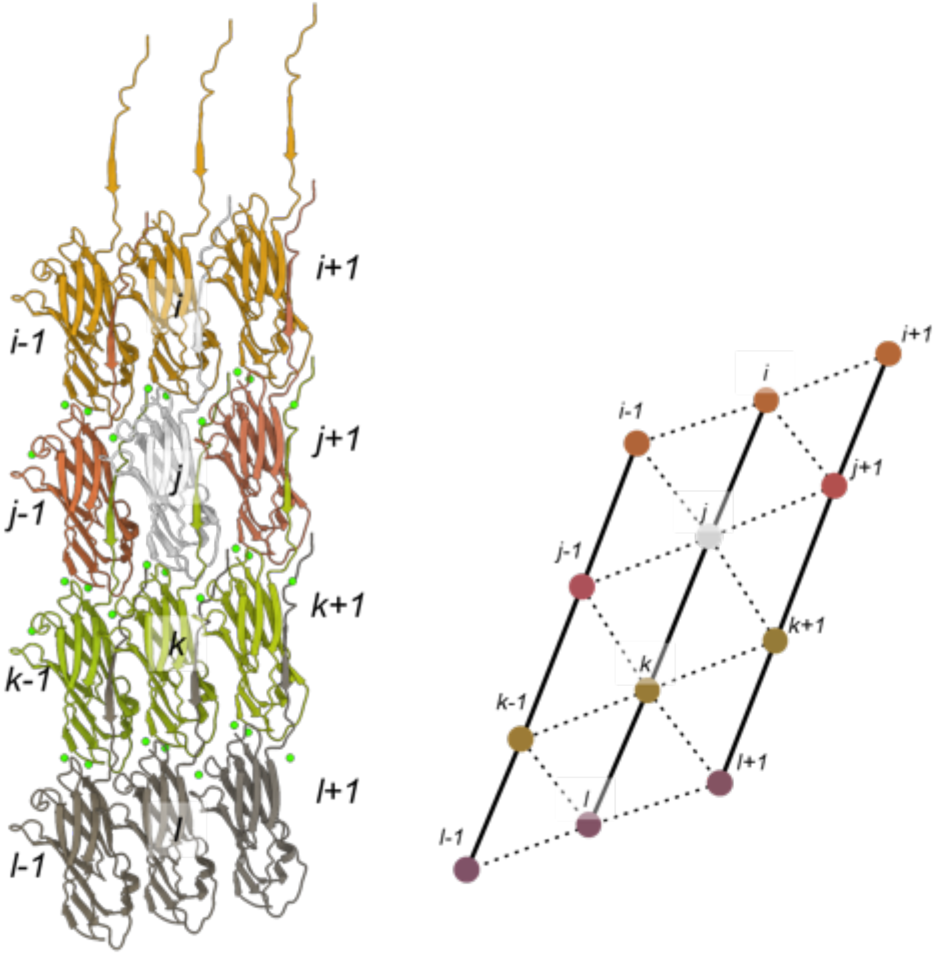


**Supplementary Figure 2: Local interaction network.** Cartoon representation of the nearest neighbor contacts with a schematic representation of the contact modes between the subunits.

```
# hmmsearch :: search profile(s) against a sequence database
# HMMER 3.0 (March 2010); http://hmmer.org/
# Copyright (C) 2010 Howard Hughes Medical Institute.
# Freely distributed under the GNU General Public License (GPLv3).
# - - - - - - - - - - - - - - - - - - - - - - - - - - - - - - - - - - - -
# query HMM file:                  canA.hmm
# target sequence database:        NZ_AP028907.1.fasta
# - - - - - - - - - - - - - - - - - - - - - - - - - - - - - - - - - - - -

Query:       canA  [M=178]
Scores for complete sequences (score includes all domains):
   --- full sequence ---   --- best 1 domain ---    -#dom-
    E-value  score  bias    E-value  score  bias    exp  N  Sequence                                  Description
    ------- ------ -----    ------- ------ -----   ---- --  --------                                  -----------
    1.5e-95  315.3   2.2    1.6e-95  315.2   1.6    1.0  1  lcl|NZ_AP028907.1_prot_WP_338250752.1_44   [locus_tag=AAA988_RS00220] [db_x
    1.3e-91  302.4   1.8    1.6e-91  302.2   1.2    1.0  1  lcl|NZ_AP028907.1_prot_WP_338251966.1_680  [locus_tag=AAA988_RS03505] [db_x
    4.4e-91  300.7   5.3    4.9e-91  300.6   3.7    1.0  1  lcl|NZ_AP028907.1_prot_WP_338251948.1_671  [locus_tag=AAA988_RS03460] [db_x
    3.6e-90  297.7   3.2      4e-90  297.6   2.2    1.0  1  lcl|NZ_AP028907.1_prot_WP_338249530.1_1835 [locus_tag=AAA988_RS09345] [db_x
    4.9e-87  287.5   3.1    5.5e-87  287.4   2.1    1.0  1  lcl|NZ_AP028907.1_prot_WP_338250768.1_52   [locus_tag=AAA988_RS00260] [db_x
    6.5e-86  283.9   3.6    7.2e-86  283.7   2.5    1.0  1  lcl|NZ_AP028907.1_prot_WP_338248767.1_1455 [locus_tag=AAA988_RS07430] [db_x
    1.7e-75  249.9   0.5    1.9e-75  249.7   0.3    1.0  1  lcl|NZ_AP028907.1_prot_WP_338250730.1_31   [locus_tag=AAA988_RS00155] [db_x
    9.1e-58  192.2   1.2      1e-57  192.0   0.8    1.0  1  lcl|NZ_AP028907.1_prot_WP_338251989.1_691  [locus_tag=AAA988_RS03560] [db_x
    9.9e-58  192.1   1.8    1.2e-57  191.8   1.2    1.1  1  lcl|NZ_AP028907.1_prot_WP_338248772.1_1459 [locus_tag=AAA988_RS07450] [db_x
    3.6e-42  141.4   0.0    3.6e-32  108.8   0.0    2.2  2  lcl|NZ_AP028907.1_prot_WP_338251974.1_684  [locus_tag=AAA988_RS03525] [db_x
      3e-31  105.8   0.7    7.6e-28   94.7   0.0    2.1  2  lcl|NZ_AP028907.1_prot_WP_338251566.1_478  [locus_tag=AAA988_RS02490] [db_x
    5.7e-29   98.4   1.0    1.4e-26   90.5   0.0    2.1  2  lcl|NZ_AP028907.1_prot_WP_338251541.1_466  [locus_tag=AAA988_RS02430] [db_x
    1.2e-28   97.3   0.0    1.7e-28   96.8   0.0    1.2  1  lcl|NZ_AP028907.1_prot_WP_338248751.1_1447 [locus_tag=AAA988_RS07390] [db_x
    1.3e-27   94.0   0.0    2.4e-27   93.1   0.0    1.4  1  lcl|NZ_AP028907.1_prot_WP_338251950.1_672  [locus_tag=AAA988_RS03465] [db_x
    2.5e-25   86.5   3.6    2.3e-24   83.3   0.4    2.1  2  lcl|NZ_AP028907.1_prot_WP_338251550.1_471  [locus_tag=AAA988_RS02455] [db_x
      8e-21   71.8   0.0    1.3e-09   35.3   0.0    2.3  2  lcl|NZ_AP028907.1_prot_WP_338251545.1_468  [locus_tag=AAA988_RS02440] [db_x
    5.2e-16   56.2   5.8      2e-14   51.0   4.0    2.0  1  lcl|NZ_AP028907.1_prot_WP_338251946.1_670  [locus_tag=AAA988_RS03455] [db_x
    5.4e-09   33.3   0.0    9.6e-09   32.5   0.0    1.3  1  lcl|NZ_AP028907.1_prot_WP_338248774.1_1460 [locus_tag=AAA988_RS07455] [db_x
    5.7e-07   26.7   0.0    9.2e-07   26.0   0.0    1.3  1  lcl|NZ_AP028907.1_prot_WP_338251995.1_694  [locus_tag=AAA988_RS03575] [db_x
```

**Supplementary Figure 3: Search for CanA-like subunits in the *Pyrodictium abyssi* genome.** Output of a HMMer 3.0 search against NZ_AP028907.1 using CanA (UniParc: UPI0024181436) as query sequence.

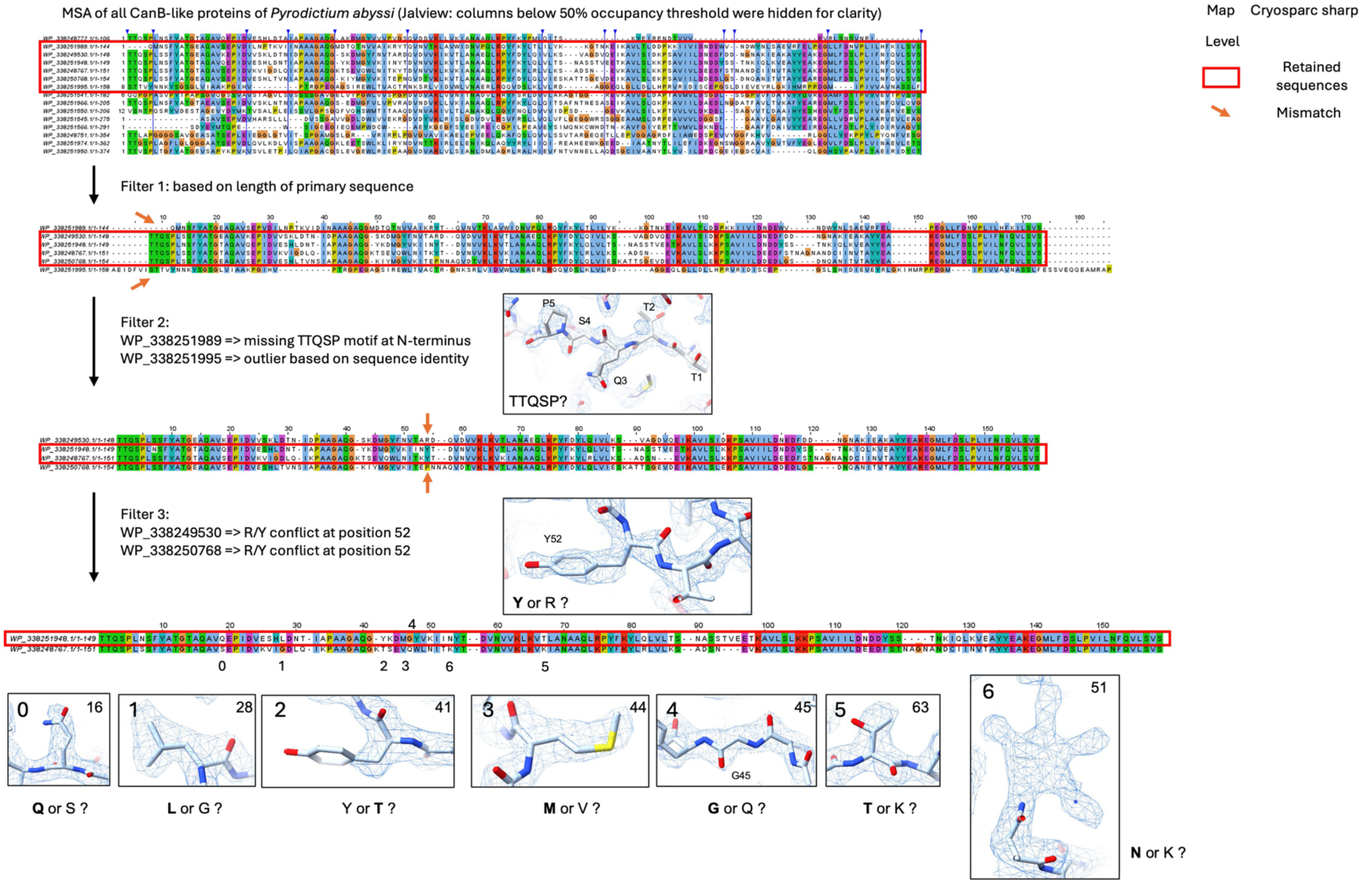


**Supplementary Figure 4: Identification of the major subunit of *ex vivo* cannulae from *Pyrodictium abyssi*.** Multiple sequence alignment (MSA) of all the putative cannulae subunits that were identified using a HMMer search against the *Pyrodictium abyssi* AV2 genome (NZ_AP028907.1). MSA was constructed in Jalview 2.11.3.3 using Muscle with preset “Protein alignment”. Subunit identification proceeded through a series of steps based on sequence length criteria and (mis)matches between amino acid side chain type and the cryoEM volume. WP_338251948.1 (annotated as “hypothetical protein”, InterProScan: no identified Pfam or Panther domains) was retained as the major subunit. Given the similarity (49.0% sequence identity) to previously identified CanA (UPI0024181436), and the fact that this sequence was derived from an *ex vivo* analysis, we propose “CanX” as a naming convention.

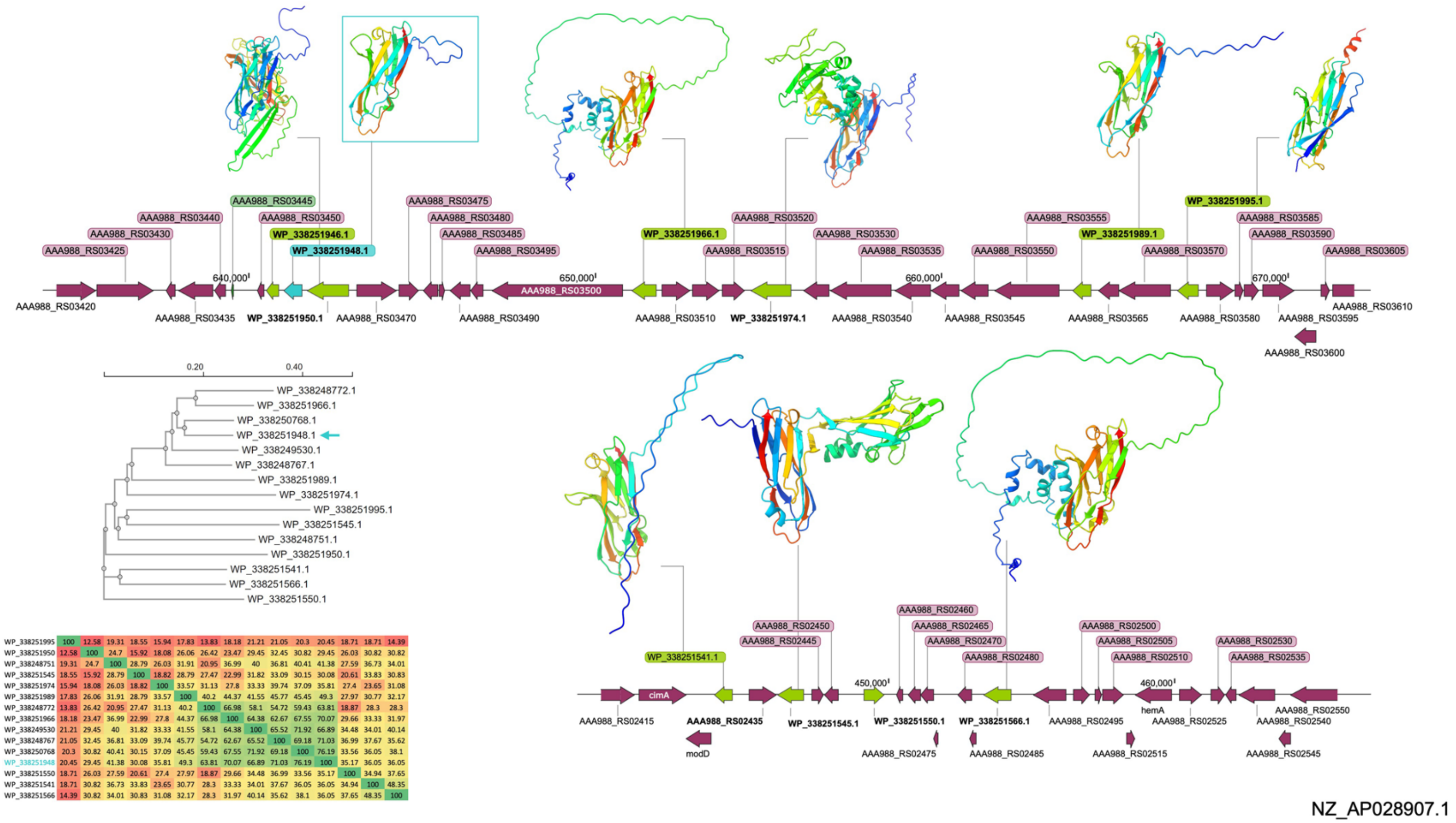


| | | | | | | | | | | | | | | | |
|---|---|---|---|---|---|---|---|---|---|---|---|---|---|---|---|
| WP_338251995 | 100 | 12.58 | 19.31 | 18.55 | 15.94 | 17.83 | 13.83 | 18.18 | 21.21 | 21.05 | 20.3 | 20.45 | 18.71 | 18.71 | 14.39 |
| WP_338251950 | 12.58 | 100 | 24.7 | 15.92 | 18.08 | 26.06 | 26.42 | 23.47 | 29.45 | 32.45 | 30.82 | 29.45 | 26.03 | 30.82 | 30.82 |
| WP_338248751 | 19.31 | 24.7 | 100 | 28.79 | 26.03 | 31.91 | 20.95 | 36.99 | 40 | 36.81 | 40.41 | 41.38 | 27.59 | 36.73 | 34.01 |
| WP_338251545 | 18.55 | 15.92 | 28.79 | 100 | 18.82 | 28.79 | 27.47 | 22.99 | 31.82 | 33.09 | 30.15 | 30.08 | 20.61 | 33.83 | 30.83 |
| WP_338251974 | 15.94 | 18.08 | 26.03 | 18.82 | 100 | 33.57 | 31.13 | 27.8 | 33.33 | 39.74 | 37.09 | 35.81 | 27.4 | 23.65 | 31.08 |
| WP_338251989 | 17.83 | 26.06 | 31.91 | 28.79 | 33.57 | 100 | 40.2 | 44.37 | 41.55 | 45.77 | 45.45 | 49.3 | 27.97 | 30.77 | 32.17 |
| WP_338248772 | 13.83 | 26.42 | 20.95 | 27.47 | 31.13 | 40.2 | 100 | 66.98 | 58.1 | 54.72 | 59.43 | 63.81 | 18.87 | 28.3 | 28.3 |
| WP_338251966 | 18.18 | 23.47 | 36.99 | 22.99 | 27.8 | 44.37 | 66.98 | 100 | 64.38 | 62.67 | 67.55 | 70.07 | 29.66 | 33.33 | 31.97 |
| WP_338249530 | 21.21 | 29.45 | 40 | 31.82 | 33.33 | 41.55 | 58.1 | 64.38 | 100 | 65.52 | 71.92 | 66.89 | 34.48 | 34.01 | 40.14 |
| WP_338248767 | 21.05 | 32.45 | 36.81 | 33.09 | 39.74 | 45.77 | 54.72 | 62.67 | 65.52 | 100 | 69.18 | 71.03 | 36.99 | 37.67 | 35.62 |
| WP_338250768 | 20.3 | 30.82 | 40.41 | 30.15 | 37.09 | 45.45 | 59.43 | 67.55 | 71.92 | 69.18 | 100 | 76.19 | 33.56 | 36.05 | 38.1 |
| WP_338251948 | 20.45 | 29.45 | 41.38 | 30.08 | 35.81 | 49.3 | 63.81 | 70.07 | 66.89 | 71.03 | 76.19 | 100 | 35.17 | 36.05 | 36.05 |
| WP_338251550 | 18.71 | 26.03 | 27.59 | 20.61 | 27.4 | 27.97 | 18.87 | 29.66 | 34.48 | 36.99 | 33.56 | 35.17 | 100 | 34.94 | 37.65 |
| WP_338251541 | 18.71 | 30.82 | 36.73 | 33.83 | 23.65 | 30.77 | 28.3 | 33.33 | 34.01 | 37.67 | 36.05 | 36.05 | 34.94 | 100 | 48.35 |
| WP_338251566 | 14.39 | 30.82 | 34.01 | 30.83 | 31.08 | 32.17 | 28.3 | 31.97 | 40.14 | 35.62 | 38.1 | 36.05 | 37.65 | 48.35 | 100 |

**Supplementary Figure 5: Genetic organization of the cannulae-like genes:** (a) putative cannulae subunits were identified in the GCF_036323395.1assembly through protein homology search (HMMer) and mapped onto the *P. abyssi* AV2 genome (NZ_AP028907.1). ORFs corresponding to cannula-like subunits are shown in green, and CanX (WP_338251948.1) is shown in cyan. AlphaFold2 models were generated using *localcolabfold* with default settings; (b) The phylogenetic tree and (c) the percent identity matrix was generated using muscle (https://www.ebi.ac.uk/jdispatcher/msa/muscle; ClustalW).

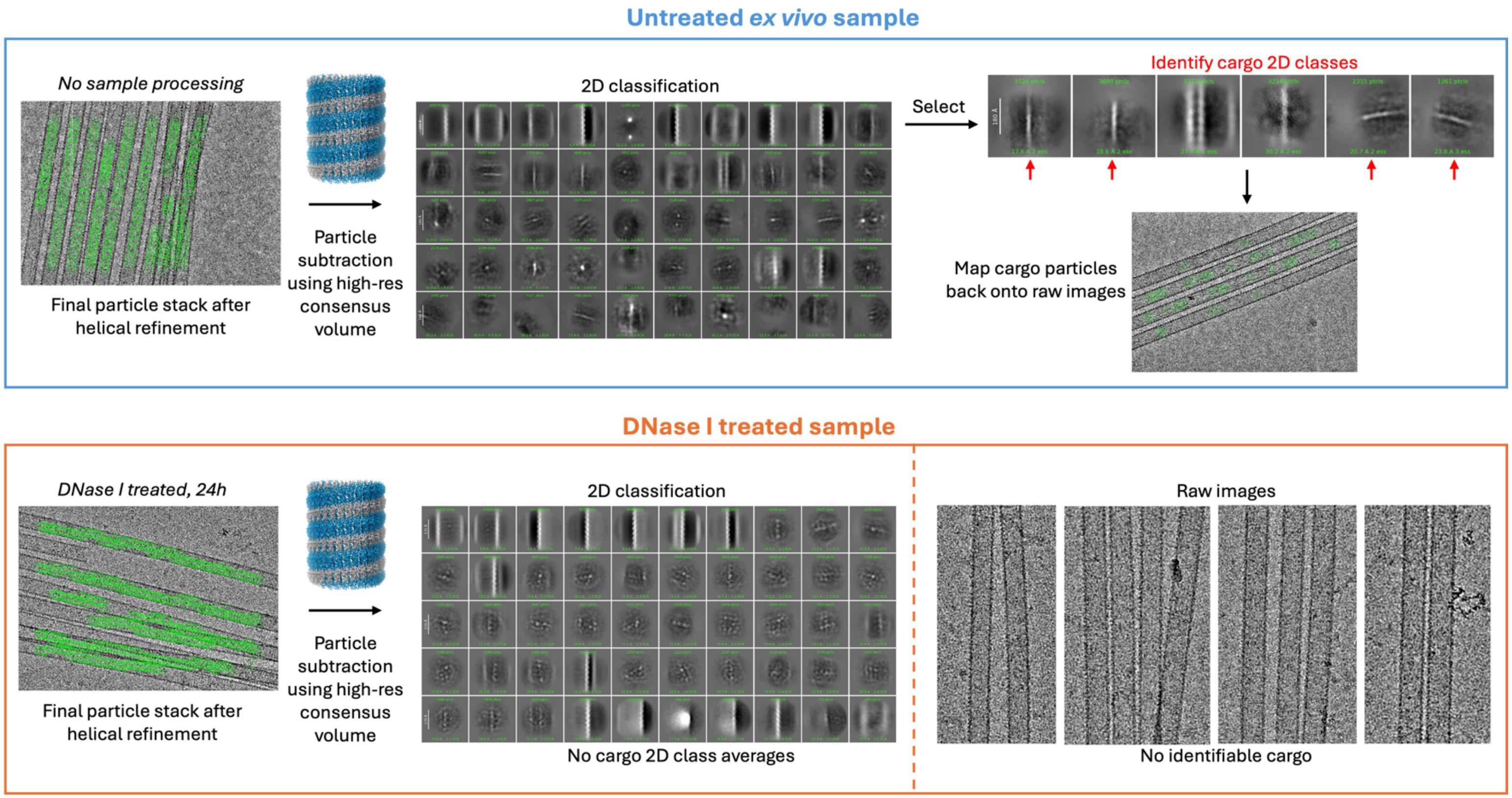


**Supplementary Figure 6: Determination of the fibril cargo identity:** CryoEM datasets were collected and processed for both an untreated and DNAse I treated (24 h, 37°C, 10 mM $CaCl_2$ + 10 mM $MgCl_2$) sample. High-resolution reconstructions (non-treated: 2.3 Å; DNAse I treated: 2.46 Å) were used to in a particle subtraction job, followed by a round of 2D classification. For the non-treated sample, this resulted in 2D class averages of thin 2 nm fibrils that map onto cargo locations in the raw micrographs. No such subtracted fibril classes could be produced for the DNAse I treated dataset, nor could any cargo be identified in the raw micrographs of the DNAse I treated dataset.

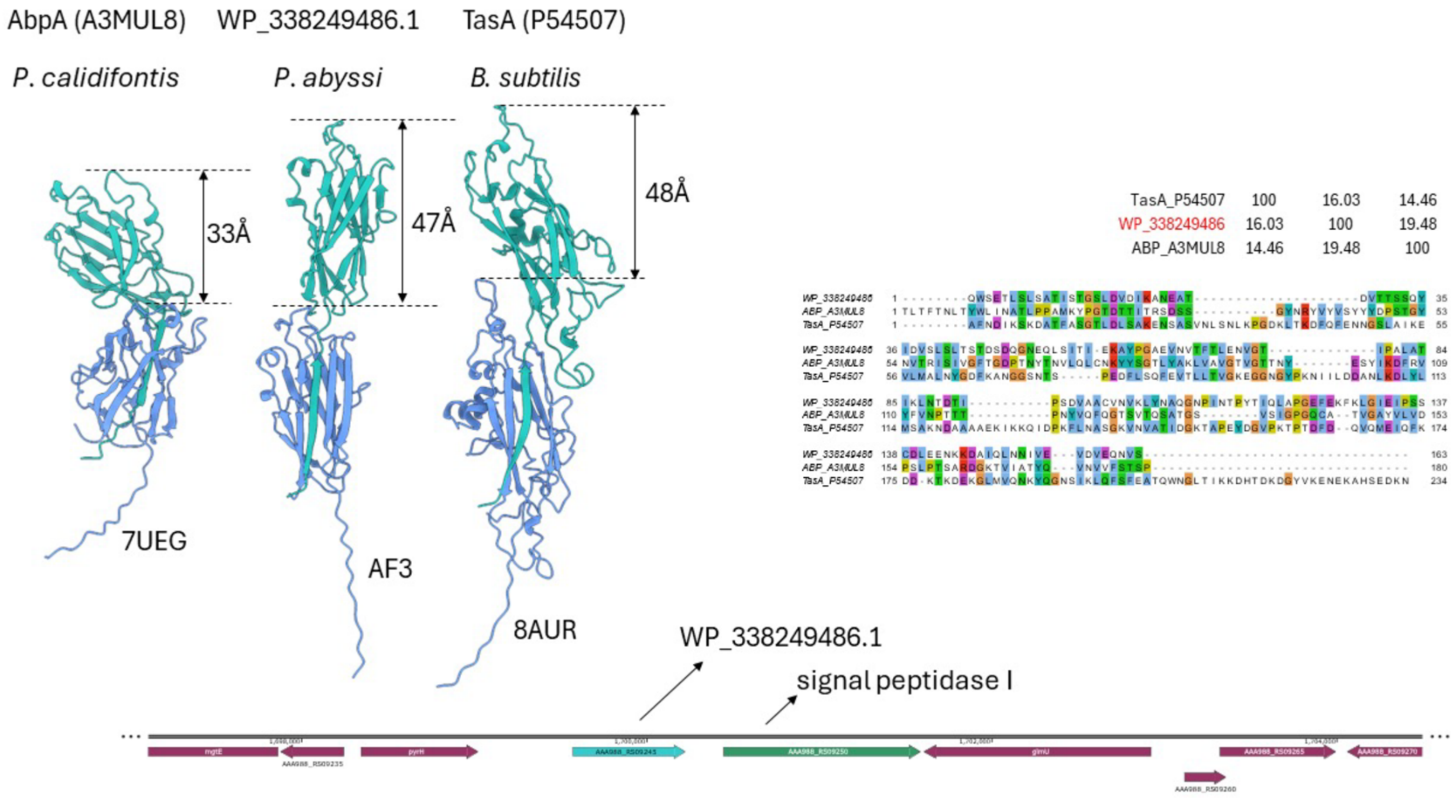


**Supplementary Figure 7:** Structural comparison between Archaeal bundling pili of *Pyrobaculum calidifontis*, extracellular matrix protein TasA from *Bacillus subtilis* and the candidate bundling pili protein (AbpX) of *Pyrodictium abyssi* AV2.

a. (M)VKYTTLAIAGIIASAAALALLAGFATTQSPLNSFYATGTAQAVSEPIDVESHLGSITPAAGAQGSDDIGYAIVWIKDQVNDVKLKVTLANAEQLKPYFKYLQIQITSGYETNSTALGNFSETKAVISLDNPSAVIVLDKEDIAVLYPDKTGYTNTSIWVPGEPDKIIVYNETKPVAILNFKAFYEAKEGMLFDSLPVIFNFQVLQVG
b. (M)KKSSSGAGSLAPLAALAASAIALALFTGYASVTSDRTYYGYGDVSVKNEPVNVVVSPFSFPGANASLSSGGQGVFKKPDWIRVVNQSDVENVKLEIDWVNANQAANYFDYARILVTGPNGQVKGYLSLQHGKAWITLDAEELREGAVLGAVMYYEVKEGVLASRLPLVFKVRVVETG

**Supplementary Figure 8: Amino acid sequences of CanA and Hyper2 that served as the basis for the recombinant proteins analyzed in this study.** (a). Full amino acid sequence of *P. abyssi* TAG11 CanA (UPI0024181436).[1] The underlined region corresponds to the ten amino acids that were deleted in the recombinant truncation of CanA, K1-CanA, which was not competent for polymerization into cannula-like filaments. (b) Full amino acid sequence of *Hyperthermus* species CanA-like protein Hyper2 (UniProt: A0A432R7L7). For all proteins, the secretion signal sequence is highlighted in yellow and the sites of putative N-glycosylation sequons (NxS/T, x ≠ P) are highlighted in cyan. Recombinant full-length CanA and Hyper2, corresponding to the respective signal-sequence processed proteins, were expressed as transcription fusions downstream of an initiation codon (AUG). The maximum likelihood positions for secretion signal peptidase cleavage were determined from SignalP 5.0 analysis.[2]

>CanA
ATGACGACGCAAAGCCCACTGAATAGCTTCTACGCAACCGGCACCGCTCAAGCCGTTAGCGAGCCGATTGACGTTGAGTCCCACCTGGGTTCG
ATCACGCCGGCGGCGGGTGCGCAAGGTTCTGACGACATTGGTTATGCAATCGTCTGGATTAAAGATCAAGTGAATGACGTGAAGTTGAAAGTAAC
CCTGGCGAACGCAGAACAGCTGAAGCCGTACTTTAAGTATCTGCAGATCCAGATTACGAGCGGCTATGAGACTAACAGCACGGCACTGGGTAAT
TTCAGCGAAACCAAAGCCGTGATCAGCCTGGACAATCCTTCCGCGGTTATCGTCCTGGATAAGGAAGATATTGCGGTTTTGTACCCGGATAAGAC
CGGTTACACCAATACCAGCATCTGGGTTCCGGGTGAGCCGGACAAAATCATTGTCTATAACGAAACCAAACCGGTGGCGATTCTGAACTTCAAG
GCTTTTTACGAAGCCAAAGAGGGCATGCTGTTTGATAGCCTGCCGGTGATCTTTAACTTCCAGGTCTTACAGGTTGGCTAATAA

>Hyper1
ATGACTGTTAACTCTGATCGTTCCTACTATGCTTATGGCGATATTACCGCGGACAATGAAAGCGTGAATATTCTGATCACGAATCCGGAGCCGAAG
ATCAGCGCGGGTGGCCAAGGTGTCCTGAATCTGTTTTGGGCAAACATCAGCAAGGGTAACGACGTCGAAAATGTCAAACTGATGTTCCGCATCG
TGAACGTTTGGCAGTTGCGTGCCTATTTGGACTACCTGAAAATTGTCCTGAATGACACGAGCAACCAAGTGAAGGGTGTTATTACCCTGCGCCAC
CCGGTTGATATTGTGACCCTGGACAACGATGACTTTAACCGTGATGGTTGGTGCAACATCAGCGCGACGGCATACTACGAAGCGAAAGAGGGCA
TGCTGTTCAGCAATCTGCCGGTGATCGTTCAGGTAAAACTGCTGCAGACCAGCTAA

>Hyper2
ATGTCTGTTACTTCTGATCGTACCTATTACGGCTATGGCGATGTTTCCGTCAAAAATGAACCTGTGAATGTTGTGGTTAGCCCGTTCAGCTTTCCAGG
TGCGAATGCAAGCCTGAGCTCGGGTGGCCAGGGTGTCTTTAAGAAACCGGACTGGATCCGTGTCGTAAACCAGAGCGATGTGGAAAACGTGAA
GTTAGAGATCGACTGGGTGAACGCGAACCAAGCTGCGAACTACTTCGACTACGCACGCATTTTGGTGACCGGTCCGAATGGCCAAGTCAAGGG
CTACCTGAGCCTGCAGCACGGTAAGGCATGGATTACGCTGGACGCGGAAGAACTGCGTGAGGGTGCCGTCTTGGGTGCCGTCATGTACTATGA
AGTTAAAGAGGGCGTGCTGGCGAGCCGCCTGCCGCTGGTTTTCAAAGTGCGTGTTGTTGAAACCGGTTAA

**Supplementary Figure 9:** Codon optimized coding sequences corresponding to the recombinant open reading frames of putative signal sequence cleaved cannula-mimetic proteins; CanA (UPI0024181436), Hyper1 (DSY37_00495, UniProt: A0A432RA20) and Hyper2 (DSY37_03180, UniProt: A0A432R7L7).

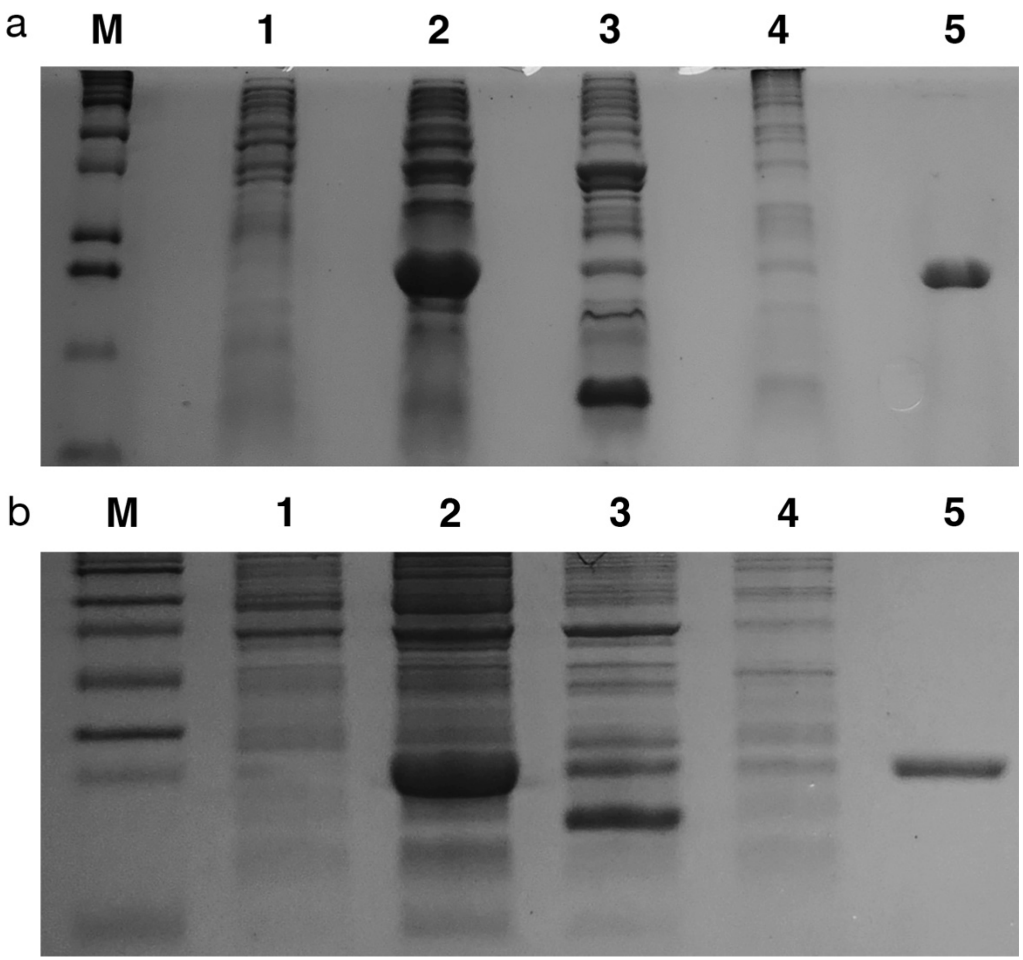


**Supplementary Figure 10: SDS-PAGE analysis of the bacterial expression and purification of cannula-mimetic proteins:** (a) CanA and (b) Hyper2. Lanes: M, protein molecular weight standards; 1, pre-induction bacterial cell culture; 2, post-induction (4 h) bacterial cell culture; 3, crude cell lysate; 4, heat-treated cell pellet; 5, purified fraction containing soluble cannula-mimetic proteins.

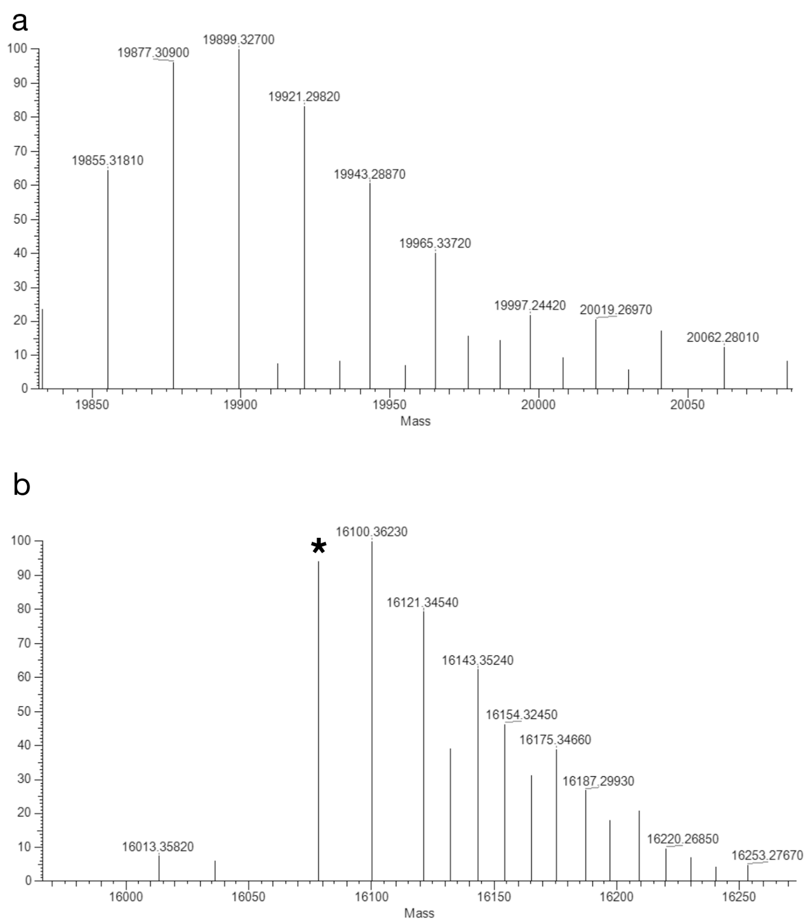


**Supplementary Figure 11: Deconvoluted electrospray-ionization mass spectra (ESI-MS) of cannula-mimetic proteins**. (a) ESI-MS of CanA. The most prominent peaks correspond to sodium ion adducts of the monoisotopic mass of CanA monomer, in which the N-terminal methionine residue has been proteolytically cleaved by methionyl aminopeptidase (calculated mass: 19,833 + 23($Na^+$) – 1($H^+$)). (b) ESI-MS of Hyper2. The most prominent peaks correspond to sodium ion adducts of the Hyper2 protein The parent ion of Hyper2 is observed at a mass of 16,078 dalton (*), which is consistent with a calculated monoisotopic mass of Hyper2 in which the N-terminal methionine residue has been proteolytically cleaved by methionyl aminopeptidase (calculated mass = 16,078 dalton). The mono-sodium ion adduct of Hyper2 monomer is observed at 16,100 dalton (calculated: 16,078 + 23($Na^+$) – 1($H^+$)).

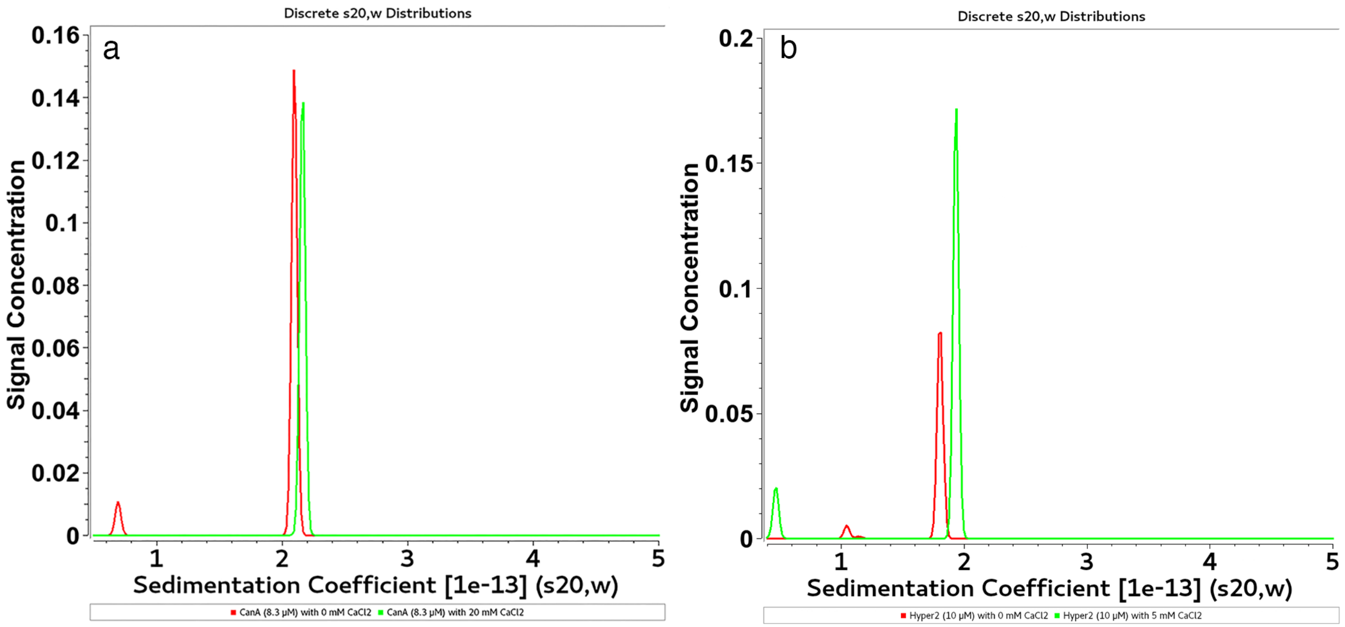


**Supplementary Figure 12: Sedimentation coefficient distribution curves determined from sedimentation velocity analytic ultracentrifugation (SV-AUC) analysis of recombinant cannula proteins.** (a) CanA (8.3 μM) (b) and Hyper2 (10 μM) in low-salt polymerization buffer (50 mM Tris-HCl pH 7.5, 80 mM NaCl, 9% glycerol, 0.01 mM EDTA) in the absence and presence of the indicated concentrations of calcium ions. *S*-values of were calculated for CanA in the absence and presence of calcium ion as 2.12S and 2.31S, which corresponded to molecular weights of 19,653 Da and 20,598 Da, respectively. The molecular weight of CanA was calculated to be 19,833 Da. *S*-values were calculated for Hyper2 in the absence and presence of calcium ion as 1.76S and 1.91S, which corresponded to molecular weights of 16,108 Da and 16,540 Da, respectively. The molecular weight of Hyper2 was calculated to be 16,078 Da.

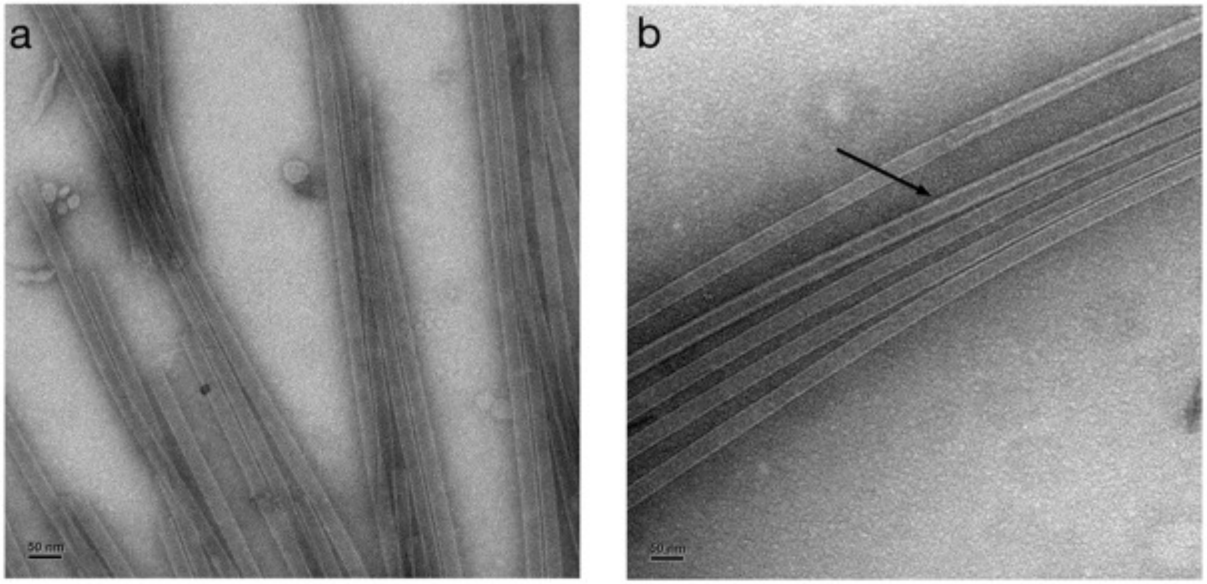


**Supplementary Figure 13**. **Representative negative-stain TEM images of cannula-mimetic protein filaments derived from calcium-ion induced self-assembly of recombinant cannula-like proteins**. (a) CanA (50 μM) after addition of $CaCl_2$ (20 mM) and thermolysis for 20 min at 80 °C. (b) Hyper2 (100 μM) after addition of $CaCl_2$ (10 mM) and incubation at ambient temperature for 24 h. Arrow indicates the presence of a thicker (multi-walled) tubular filament. Assembly experiments were performed in low-salt polymerization buffer (50 mM Tris-HCl pH 7.5, 80 mM NaCl, 9% glycerol, 0.01 mM EDTA). Scale bar: 50 nm.

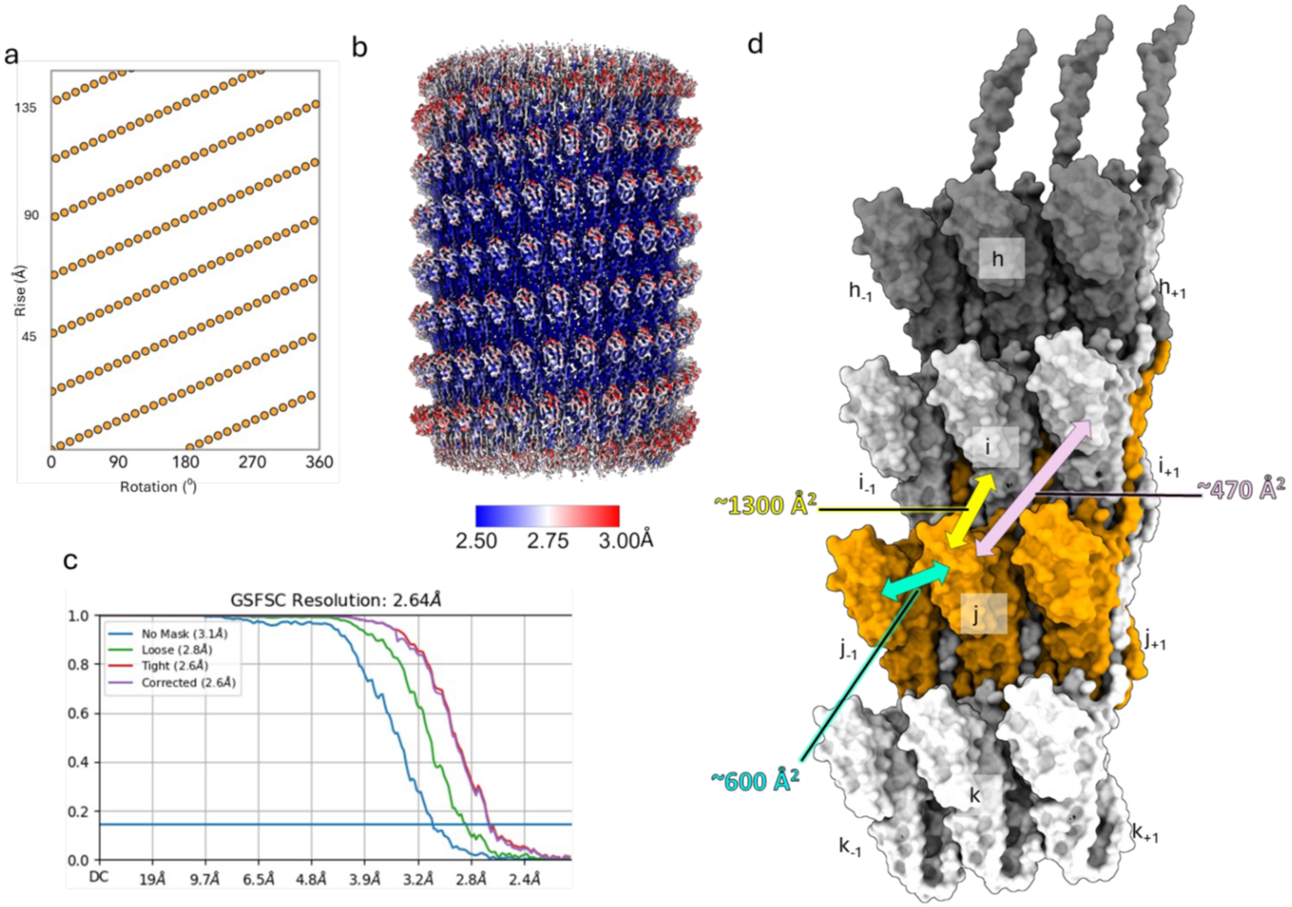


**Supplementary Figure 14: Supplementary figures to the main Figure 2 about cryoEM structure of recombinant CanA tube.** (a) Helical net, (b) Local resolution map and, (c) Map-to-map Fourier Shell Correlation (FSC) curve for cryoEM reconstruction of CanA filament. (d) Interface area between neighboring protomers calculated by PISA server.

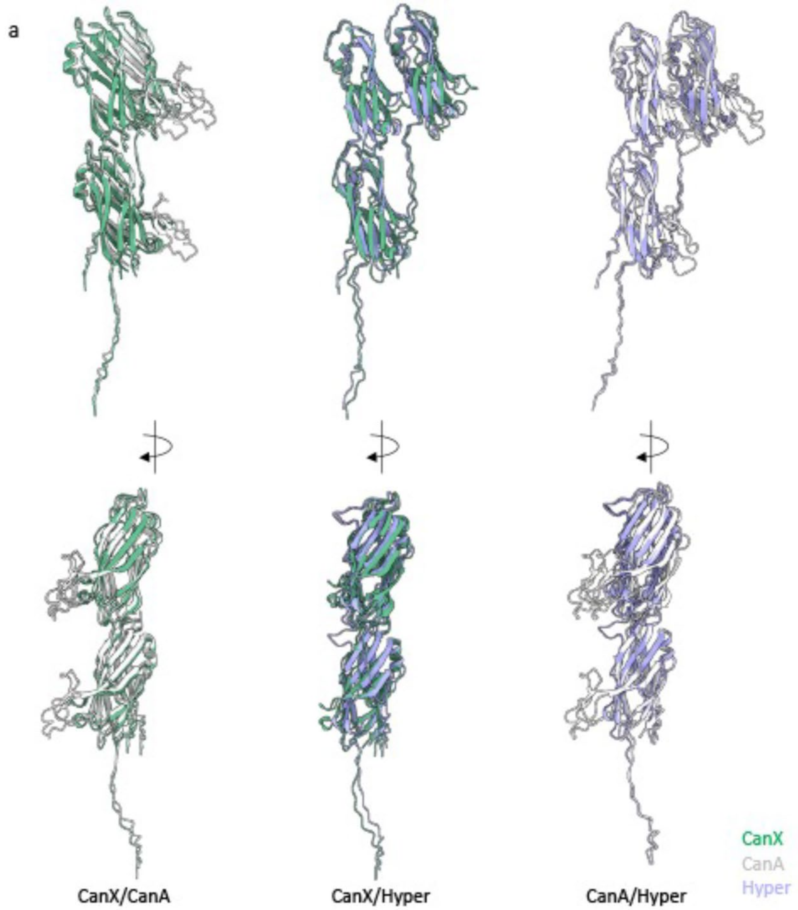


b

| *Interface* | *Protomer* | *Dimer (radial)* | *Dimer (axial)* |
|---|---|---|---|
| CanX/CanA | 0.71Å | 0.84Å | 1.0Å |
| CanX/Hyper | 0.99Å | 1.2Å | 1.2Å |
| CanA/Hyper | 0.87Å | 1.0Å | 1.0Å |

c

| Seq.Id. (%) | CanX | CanA | Hyper2 |
|---|---|---|---|
| CanX | 100 | 68.92 | 30.50 |
| CanA | 68.92 | 100 | 32.62 |
| Hyper | 30.50 | 32.62 | 100 |

**Supplementary Figure 15. Structural comparison of inter-protomer interfaces with cannula-like filaments derived from CanX (WP_338251948.1), CanA (UPI0024181436), Hyper2 (DSY37_03180, UniProt A0A432R7L7).** (a) Pairwise backbone alignments of two protomers in the respective cannula-like assemblies. (b) Calculated RMSD values for backbone alignments of individual protomers and radial (intra-protofilament) and axial (inter-protofilament interfaces between dimeric pairs of protomers within the respective cannula-like filaments. (c) Pairwise sequence identity matrix for cannula-like proteins

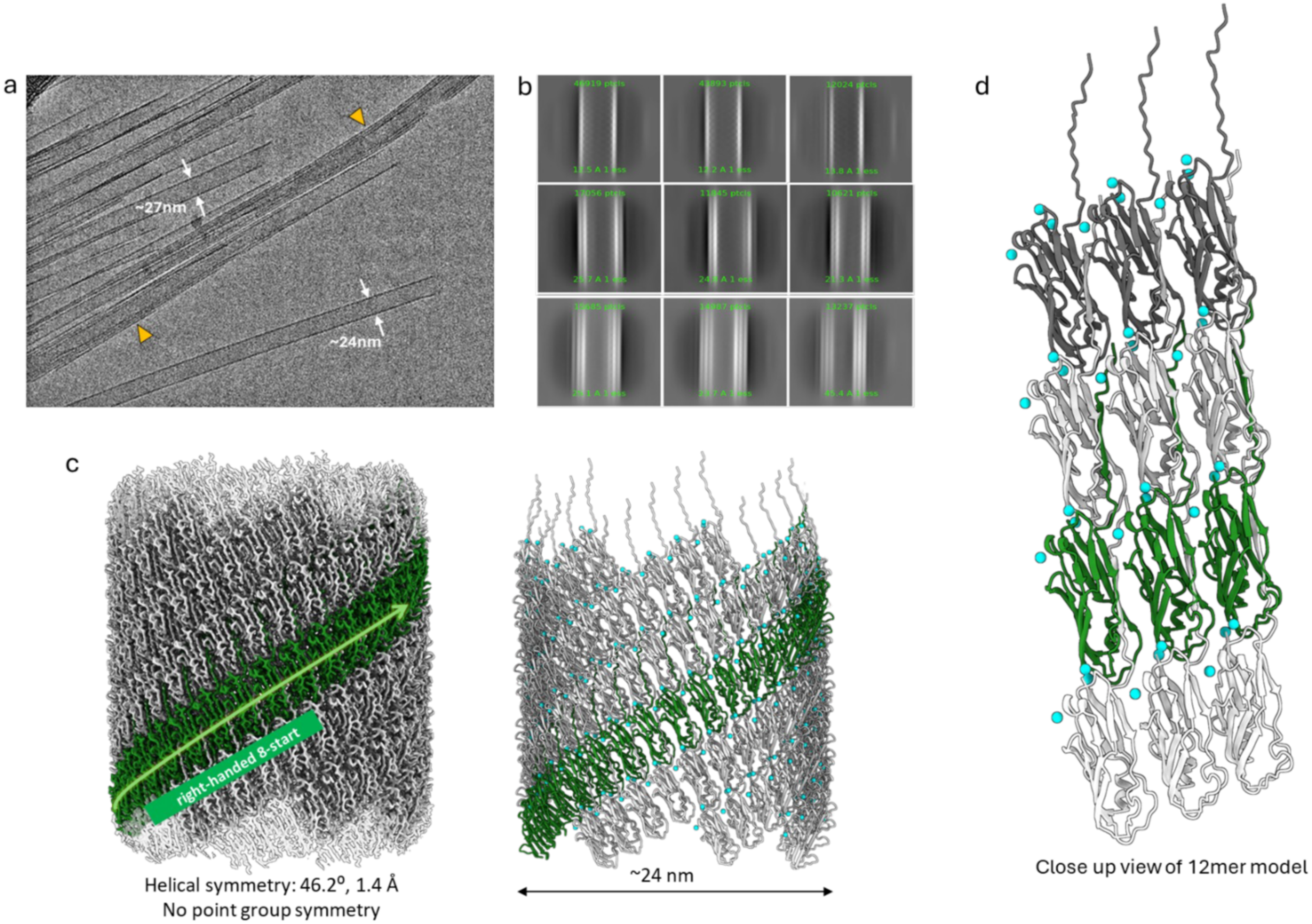


**Supplementary Figure 16: Polymorphisms in helical packing of cannula-like tubules formed by the self-assembly of the recombinant Hyper2.** (a) Representative cryoEM micrograph showing the heterogenous population of Hyper2 tubes. Multi-wall tubes are marked by orange arrowhead. (b) 2D class averages of Hyper2 tubes of different morphology. (c) CryoEM structure (map and model) of Hyper2 thin tube having different diameter and helical parameters than the Hyper2 thicker tube that is described in the main Figure 2. (d) Close-up of 12mer showing $Ca^{+2}$ ions mediated helical assembly of Hyper2 thin tube, virtually identical to the Hyper2 thicker tube.

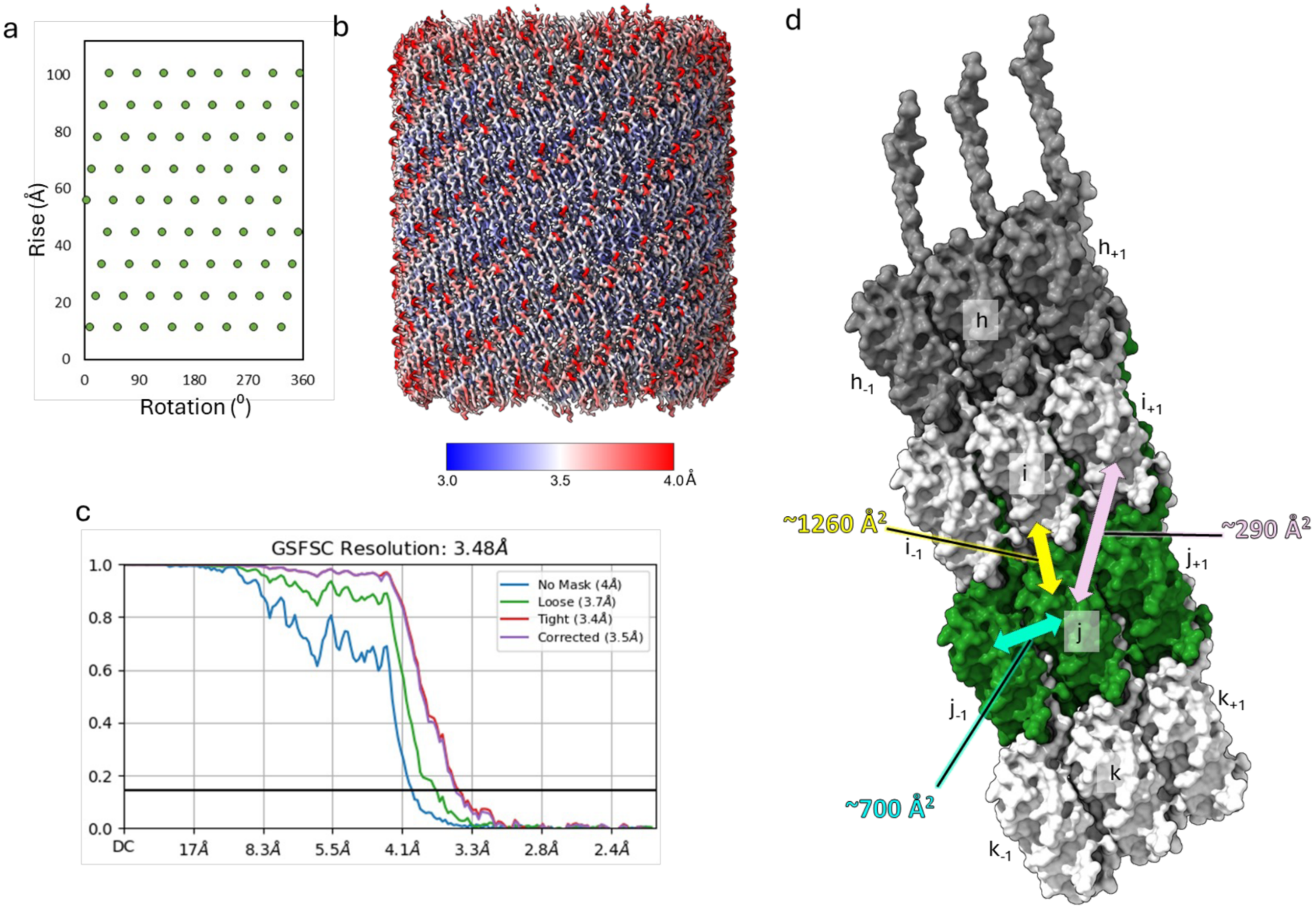


**Supplementary Figure 17. CryoEM structural analysis of *in vitro* cannula-like filament formed by the self-assembly of recombinant CanA.** (a) Helical net, (b) local resolution map and, (c) Map-to-map Fourier Shell Correlation (FSC) curve for cryoEM reconstruction of Hyper2 tube. (d) Interface area between neighboring protomers calculated by PISA server.

#### *Vulcanisaeta distributa* DSM 14429

**CanXL_VULDI (WP_054853146.1)**

MKTWLLIALAFLAGFSGMAVAANMNGVLAYLSYHLILPPPPTYSTVAAYIDLGNLTPGMSGTAEATAYLY
VNSSGYFKVELHHEGLDDVFSNFTVVVAIGNQTLTFNLHRDEYTVYLTPGNYTVSITVYYTVKPNPKGPP
NVAHKPLITIKPVGEHEEKSEGNYTGSINQGNDNDQGSED

#### *Saccharolobus solfataricus* SULM

**CanXL1_SACSO (WP_009990924.1)**

MNKLLLLGVLLSTILVGGVVIGEEISGSLGTISYNVTSPTIQTTLASFNLGNLTAGQKGTLTENATLTVS
ANGTYTIKLKEDVLGDVFSEFNVTIHIANYTFTLSLHGTEHYTLYLSKGTYTVNIKISYVVSQNPKAKSV
SNIPFLMVKYGGEQQAQQTQTAEHGKKHHDDDNEQDD

**CanXL2_SACSO (WP_009991716.1)**

MKTSILALTLVGAFLAGLATAGVAGYGPLAYISYHIMVSQQKGQAQVIPAYINLGNLTAGEKGNITATAA
INISTNGTYTIDLLHADKLQKIFSNFTVELKINNYTIILTLENDSAHVSLVNGSYNVSILIIYQVSQHPS
GDLNVNNEPLLIIHPQGEDNNS

**CanXL3_SACSO (WP_029552507.1)**

MLYFDVNASSKVYCFTVSWNQGLYGIFNQLQIVDRQTLLVGLVIITLIVNVILGVELYYTLHNRPTQPVG
GIPINVSTVQIISPSSSTTTNTTSGATSTITTTQSSTTNSGSSTTTKQPQTYSFSFKAEFKIHVKEHGIY
IVGIKPNITFSQLYVILYFEDGQVVTLNLNQTYVNVTIKENDVKVTAYIYGKSYENLTPQQILNDIGLYF
KFIGNSTNSNSNENSDEFILVRDINLVKKLFEIQPVS

#### *Metallosphaera yellowstonensis* MK1

**CanXL1_METAY (WP_009071828.1)**

MKAAYLALVALLVGSTAVAASSVAGYGPLAYITYHIINTNEGNITIVPANINLGNLTPGEKGNVTVNASV
TLSKTDNYTIMLLHLEKLKKDFSEFKAIINIGNKTITIDLDHPFAVLQLSNGTYQVHITIVYQVSQNPSG
DLNVNNEPLLIIHPGVVHKDDHHEHHGHHHDNGNDDQGDDDRGDG

**CanXL2_METAY (WP_009072785.1)**

MNKMALILLAAVVATGAAAGAANFSTLATITYTISQVTSQSSAQPQVILNTANLNLGNLTPGQLGTASAN
TTMTVNTSGNYTFELNENPIESVFSSFNVTIIVGNKTLYLTEGQNDVHTYLSNGTYQVQIKVSYQVSQNP
EGNGVQDKPLIIVKPYQGDNSPDMNSNTRSGVTSASDNSPDHSTNSSQGDNNSGSSDSGYGSDN

**CanXL3_METAY (WP_009072754.1)**

MKQNLLVGLLVLTLAVNVTLGYELYSLNHASSAHAVAPVVFENVTVYPGQNTTAEGNLTLHPFYYKYGVL
LKFPQKGEYLVGINGTVFPGEAILVQLEDGQSVLLNQNVTQGKVIVNDNVIHATIIISGLYSGSAPSPQL
VLQDVGLFFQFLRPLQVVNSSSEGGQQEIQNVTHDFDHGSVSDNDLNSSATNSTINSNTTINQDNITNQN
STSQPHDDNVTLQPHNENDTGQGTSQGSDSSQNSNSSSNSSQNAHESSSGDSNSSDSSGSGSSSSGSGSS
SGSGSSSSGSGSSSSGQGDSGDSGSGSDSSSGSGSQDN

**CanXL4_METAY (WP_009071324.1)**

MRNSVILILLVLSLGATAFSSVKLITSTITPSHPSPTAFSYNVSETLVVDKEGYYTVVIHKGNLNLNSLY
VIVVFQPLPGRGNEKTVVLTLSSESEEVKLHRGVYEVIVYFTGTTPSYEGLSMIQGNLSVTIFYSREN

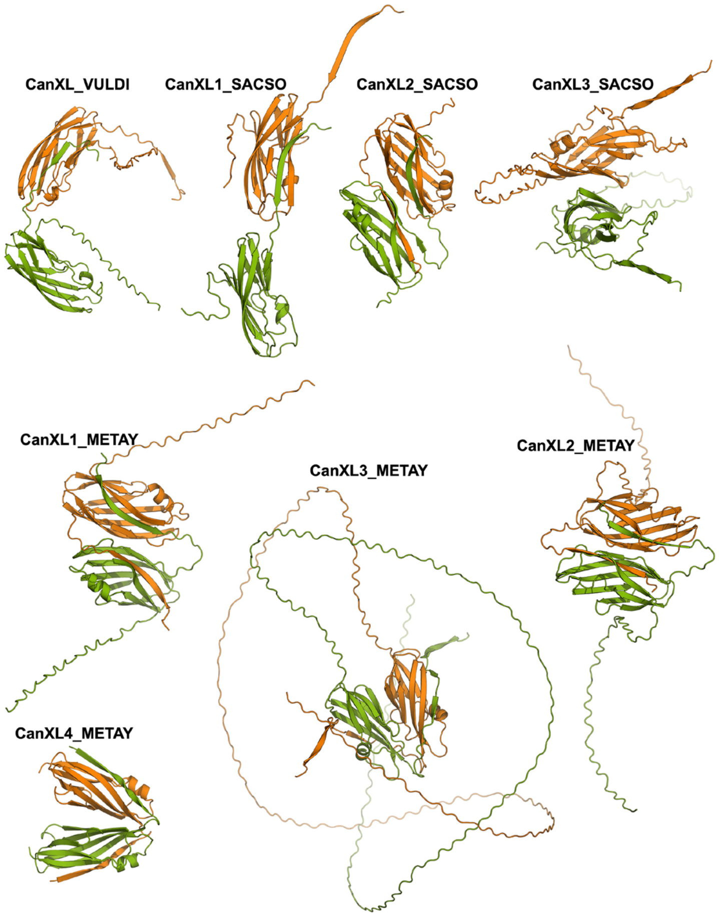


**Supplementary Figure 18. AlphaFold3 models of CanX-like proteins and their sequence features.** (a) Amino acid sequences of representative CanX-like proteins from *Vulcanisaeta distributa* DSM 14429, *Saccharolobus solfataricus* SULM, and *Metallosphaera yellowstonensis* MK1. Signal peptides and intrinsically disordered regions at the C-terminal are underlined. CanXL3_SACSO lacks a predicted signal peptide but contains a predicted transmembrane helix, highlighted in red. (b) AlphaFold3 models of dimers for the representative CanX-like proteins. All proteins, except CanXL3, dimerize by donor strand complementation. In these structures, one monomer is colored green, and the second monomer colored orange.

**Supplementary Table 1. Homologous cannula-mimetic proteins recovered from a Foldseek[3] search in which the full-length sequence of CanA was employed as a query.[1]**

| Organism | UniProt ID | Protein Length[2] | Signal Sequence Cleavage site[3,4] | Putative N-Glycosylation Sites (NxS/T) |
|---|---|---|---|---|
| *Pyrodictium occultum* | A0A0V8RWX1 | 169 aa | GFA-TT (25/26) | 1 |
| *Pyrodictium occultum* | A0A0V8RU29 | 174 aa | GFA-TT (25/26) | 2 |
| *Pyrodictium occultum* | A0A0V8RWL5 | 386 aa | GFA-TT (27/28) | 7 |
| *Pyrodictium occultum* | A0A0V8RWQ8 | 414 aa | VFA-TT (24/25) | 4 |
| *Pyrodictium occultum* | A0A0V8RWP2 | 422 aa | GFA-TT (27/28) | 9 |
| *Hyperthermus sp.* | A0A432RA20[5] | 169 aa | AYA-TV (26/27) | 4 |
| *Hyperthermus sp.* | A0A432R7L7[6] | 178 aa | GYA-SV (31/32) | 2 |
| *Hyperthermus sp.* | A0A432R7D2 | 257 aa | ALA-SI (28/29) | 0 |
| *Hyperthermus sp.* | A0A432R7C7 | 310 aa | AYA-TV (24/25) | 3 |

[1]An additional 19 sequences containing cannula-like protein domains were recently deposited from *P. abyssi* strain AV2 but are not annotated in UniProt. One of these sequences, WP_338251948.1, was used as the basis for the construction of the atomic model for the *ex vivo* cannula structure. [2]Protein length includes the N-terminal secretion signal peptide. [3]Maximum likelihood signal sequence cleavage site was determined using SignalP 5.0[2] for archaeal organisms (https://services.healthtech.dtu.dk/services/SignalP-5.0/). [4]Maximum likelihood cleavage site is reported in terms of the flanking amino acids as well as the residue positions at which cleavage is predicted to occur in the parent protein sequence. [5]The amino acid sequence of the mature protein, i.e., signal peptide cleavage product, was employed as the basis for design of the protein Hyper1.[6]The amino acid sequence of the mature protein, i.e., signal peptide cleavage product, was employed as the basis for design of the protein Hyper2.

**Supplementary Table 2. CryoEM and Refinement Statistics of Cannula-like Filaments**

| Parameter | CanA | Hyper2 - thick tube | Hyper2 – thin tube | CanX |
|---|---|---|---|---|
| **Data collection and processing** | | | | |
| Voltage (kV) | 300 | 300 | 300 | 300 |
| Electron exposure ($e^{-}$ Å$^{-2}$) | 50 | 50 | 50 | 60 |
| Pixel size (Å) | 1.08 | 1.08 | 1.08 | 0.71 |
| Particle images (n) | 659,059 | 140,207 | 95,418 | 310,172 |
| Shift (pixel) | 13 | 24 | 24 | 35 |
| **Helical symmetry** | | | | |
| Point group | C1 | C8 | C1 | C2 |
| Helical rise (Å) | 1.59 | 11.16 | 46.23 | 3.18 |
| Helical twist (°) | -173.75 | 9.59 | 1.424 | 12.10 |
| **Map resolution (Å)** | | | | |
| Map:map FSC (0.143) | 2.6 | 3.5 | 3.5 | 2.3 |
| Model:map FSC (0.38) | 2.7 | 3.5 | 3.7 | 2.4 |
| $d_{99}$ | 2.9 | 3.2 | 3.9 | 3.3 |
| **Refinement and Model validation** | | | | |
| Ramachandran Favored (%) | 94.9 | 94.48 | 93.1 | 95.24 |
| Ramachandran Outliers (%) | 0.0 | 0.0 | 0.0 | 0.0 |
| RSCC | 0.87 | 0.85 | | 0.42 |
| Clash score | 3.9 | 5.74 | 5.31 | 6.04 |
| Bonds RMSD, length (Å) | 0.002 | 0.007 | 0.008 | 0.003 |
| Bonds RMSD, angles (°) | 0.590 | 0.741 | 0.783 | |
| **Deposition ID** | | | | |
| PDB (model) | 7UII | 9BAB | 9BAC | 9H8B |

| | | | | |
|---|---|---|---|---|
| EMDB (map) | EMD-26546 | EMD-44403 | EMD-44404 | EMD-51935 |

observations for a given reflection.
[d] $R = \sum \left| |F_o| - |F_c| \right| / \sum \left| |F_o| \right|$, where $F_o$ and $F_c$ denote observe and calculated structure factors, respectively.
[e] $R_{free}$ was calculated using 5% of data excluded from refinement.
[f] Calculated using Molprobity.

### References


1 Kreitner, R. *et al.* Complete sequential assignment and secondary structure prediction of the cannulae forming protein CanA from the hyperthermophilic archaeon Pyrodictium abyssi. *Biomol NMR Assign* **14**, 141-146 (2020). https://doi.org:10.1007/s12104-020-09934-x

2 Almagro Armenteros, J. J. *et al.* SignalP 5.0 improves signal peptide predictions using deep neural networks. *Nat Biotechnol* **37**, 420-423 (2019). https://doi.org:10.1038/s41587-019-0036-z

3 van Kempen, M. *et al.* Fast and accurate protein structure search with Foldseek. *Nat Biotechnol* **42**, 243-246 (2024). https://doi.org:10.1038/s41587-023-01773-0
